# Integration of single-cell RNA and protein data identifies novel clinically-relevant lymphocyte phenotypes in breast cancers

**DOI:** 10.1101/2022.05.31.494081

**Authors:** Ghamdan Al-Eryani, Nenad Bartonicek, Chia-Ling Chan, Alma Anderson, Kate Harvey, Sunny Z. Wu, Dan Roden, Taopeng Wang, John Reeves, Bertrand Z Yeung, Etienne Masle-Farquhar, Christopher C Goodnow, Cindy Ma, Tri G. Phan, Joakim Lundeberg, Simon Junankar, Alexander Swarbrick

## Abstract

Immune cells are critical determinants of solid tumour aetiology, but the diverse phenotypes of intra-tumoural immune cells remain incompletely characterised. We applied integrated single cell RNA sequencing (scRNA-Seq) and highly multiplexed protein epitope analysis to a cohort of breast cancer samples to resolve cell states within the tumour microenvironment. We reveal novel protein markers for resting and activated tumour infiltrating lymphocytes, and show that high expression of CD103 primarily marks exhausted CD8 rather than tissue resident CD8 T-cells in human breast cancers. We identify two distinct states of activated CD4+ T follicular helper (Tfh) cells. A population resembling conventional Tfh (cTfh) cells were localised primarily to lymphoid aggregates by spatial transcriptomics. In contrast, cancer associated Tfh (caTfh) cells expressing markers of tissue residency and exhaustion co-localized with cancer foci and signalled to macrophages. Importantly, increased caTfh : cTfh ratio associated with improved disease outcome and response to checkpoint immunotherapy.

## Introduction

Solid tumours constitute a diverse ecosystem of cells whose functions can be coordinated by malignant cells to promote their uncontrolled growth and spread (Hanahan & Weinberg, 2011). The successes of immune and stromal targeted therapies in certain cancer types and individuals but not others underscore the dynamic and variable intercellular relationships engaged by cancer cells to sustain malignancy(Bruni et al., 2020; Valkenburg et al., 2018). Comprehensive phenotyping of cell types in the tumour microenvironment (TME) is crucial to understand and manipulate tumour immunity for patient benefit. Although breast cancers are characterised by relatively low mutational load and lower immunogenicity, a role for tumour immune infiltration in disease outcome has nonetheless been shown(Ayse Bassez et al., 2021; P. Savas et al., 2016; Peter Savas et al., 2018; Wu et al., 2021; Y. Zhang et al., 2021). For example, the interplay between tumour infiltrating lymphocytes (TILs) and tumor associated macrophages (TAMs) has been proposed to stratify patient prognosis and response to chemotherapy (Cassetta et al., 2019; M. Molgora et al., 2020).

High-throughput single-cell RNA sequencing (scRNA-seq) is a powerful platform for characterising the cellular constituents of heterogeneous biological systems such as the TME (Azizi et al., 2018; Sijin Cheng et al., 2021). This method allows for the simultaneous measurement of thousands of mRNAs from thousands of single cells per sample(Sijin Cheng et al., 2021; La Manno et al., 2018; Svensson et al., 2018). However, scRNA-seq is limited by the sensitivity of RNA measurements at single-cell resolution, and RNA expression does not always correlate well with protein expression (Akan et al., 2012; Buccitelli & Selbach, 2020; Schwanhäusser et al., 2011). Cellular indexing of transcriptomes and epitopes by sequencing (CITE-Seq) combines scRNA-seq with detection of antibody-derived tags (ADT) as surrogates for cell surface protein levels (Peterson et al., 2017; Stoeckius et al., 2017). Proteogenomics via CITE-Seq permits the integration of transcriptome information with decades of immunological studies that have dissected immune subsets and activation states using protein readouts. CITE-Seq has proven useful in improving upon scRNA-seq-based cellular profiling, mostly in blood samples (Hao et al., 2021; Triana et al., 2021). Here we demonstrate how cellular proteogenomics enhances stratification of cell types within solid tumour microenvironments, and allows identification of lymphocyte subsets with prognostic associations. Using a dataset of 7515 cells from 6 breast cancer patients, we dissect TIL subsets that were indistinguishable when based on transcriptomics alone. We assign protein markers to phenotypes previously identified in scRNA-seq studies to play a vital role in tumour immunity, and highlight discrepancies with protein derived studies. We reveal patterns of RNA and protein co-expression across lymphocytes, and identify new protein markers of activated tissue resident T-cells and innate lymphoid cells (ILCs). We find that activated tumour-associated T follicular helper (Tfh) cells differentiate into two distinct states, demarcated by the expression of either *IGFL2* and *NMB* or *HAVCR2(TIM3), LAG3* and CD103 (*ITGAE)*. Spatial transcriptomics reveals differential localization of these two Tfh cell subsets within the TME and unique signalling potential with neighbouring cells. Supporting the clinical importance of precise cellular phenotyping, we show that a signature of CD103+ Tfh cells associates with improved survival in breast cancer and correlates with improved response to anti-PD-1 therapy. These data underscore the value of integrated cellular phenotyping in complex tissue environments to identify cell types and states integral to tumour biology and clinical outcome.

## Results

### Enhanced phenotyping of the tumour microenvironment through integrated RNA and Protein based clustering

To better characterize the native tumour immune microenvironment of human breast cancer, we applied a CITE-Seq panel of 97-157 antibodies (157 in 5 samples, 97 in one; **Table S1**) to 6 breast cancer samples, including at least one from each major clinical subtype: Luminal (Estrogen-positive (ER+) and Progesterone-positive/negative (PR+/-), human epidermal growth factor receptor positive (HER2+) and Triple negative breast cancer (TNBC) (**Table S2**). A total of 16,423 cells passed our quality filter threshold (Methods) (Figure S1A), with a variable number of ADT significantly enriched in each sample (MAST test; p_adj < 0.01) and a subset of 32 ADTs common to all samples (Figure S1B). Cells were partitioned using a weighted nearest neighbour (WNN) approach that integrates RNA and surface protein expression (Hao et al., 2021). A total of 52 clusters were identified through integrated clustering and each is labelled based on its most distinctive RNA and ADT (MAST test; p_adj < 0.01; Figure 1A-B). These 52 cellular phenotypes collapsed into 27 clusters using RNA alone or 16 clusters using only ADT data for clustering (Figure 1C; Figure S1D-E). Despite moderate cell numbers and without employing a lineage specific sub-clustering analysis(S. Cheng et al., 2021; Mulder et al., 2021; Wu et al., 2021; Zheng et al., 2021), this strategy allowed us to differentiate monocytes and macrophages into 3 distinct groups each: C28-Mono:*AIF1* ADT-CD32hi, C29-Mono:*FCN1* ADT-CR1 and C30-Mono:*IFI30* ADT-CD16, and C24-Macro:*CXCL10* ADT-CD69, C25-Macro:*SSP1* ADT-CD47, C26-Macro:*SELENOP* ADT-CD158e1 (Figure 1A-B), versus a single transcriptome-based cluster for each (Figure S1D). These clusters demarcate phenotypes previously found to be relevant to patient prognosis such as TREM2-high lipid associated macrophages (LAM) from CXCL10+ Macrophages (Cassetta et al., 2019; Wu et al., 2021), or inflammatory monocytes which exhibit cDC2-like profile from classical *CD16+* monocytes(S. Cheng et al., 2021; Wu et al., 2021).

**Figure 1:**
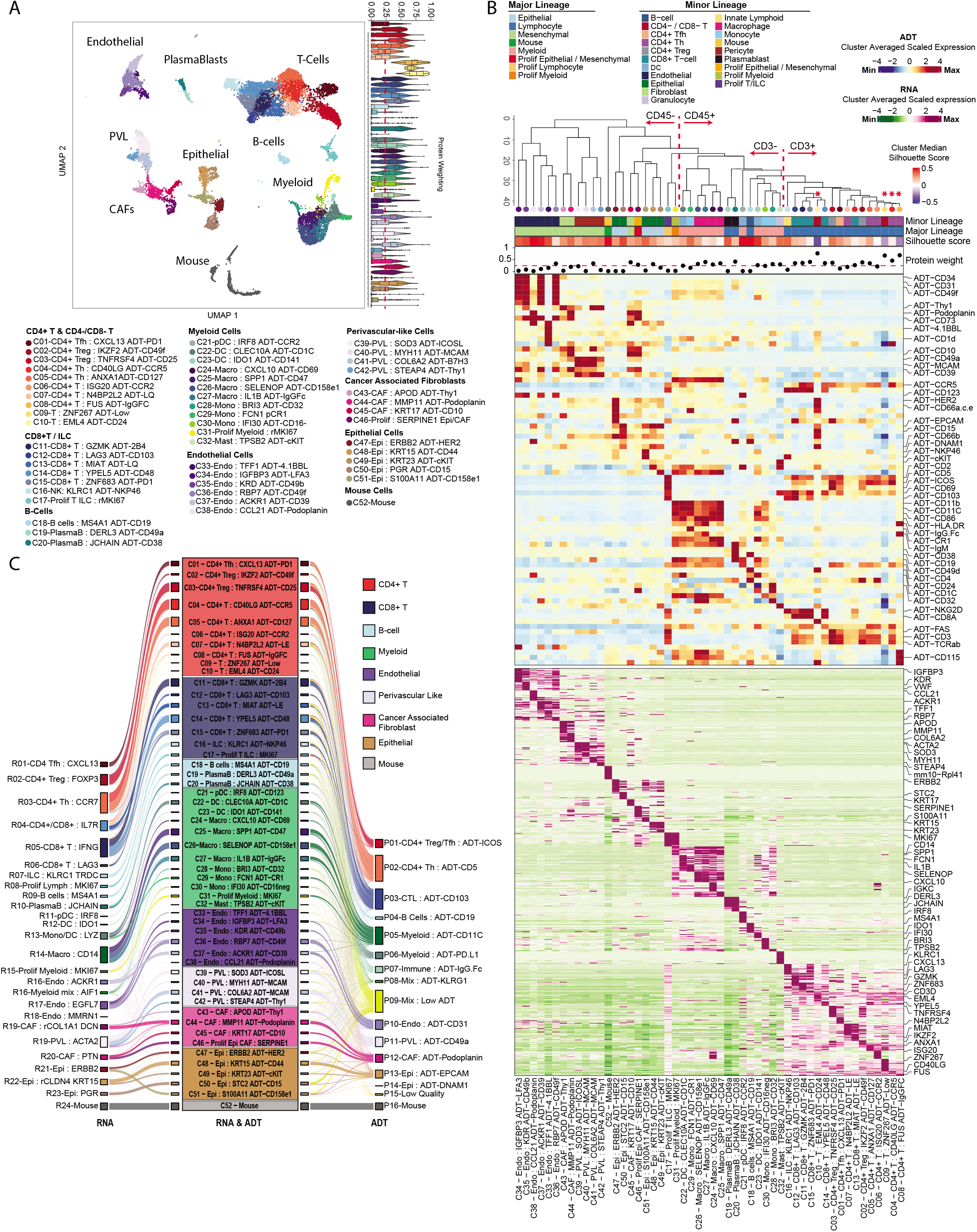
Integrated proteogenomic analysis of the breast tumour microenvironment enhances clustering resolution when compared to ADT or RNA derived clustering. **(A)** UMAP of RNA and ADT integrated clustering analysis identified 52 cell clusters within 6 human breast cancer samples. Violin plots on the right display the relative contribution of ADT markers to the definition of each cluster (“protein weighting”; (Hao et al., 2021)) and the dashed line marks the median value across all clusters (0.246). **(B)** Differentially enriched ADT (top) and RNA (bottom) features for each cluster (MAST test; p_adj < 0.05), with defining RNA and ADT features labelled on the right. Top annotation dendrogram indicates the cluster relationship derived from integrated RNA and ADT PCA values. Top bar annotations provide the mean silhouette score and median weighted protein weighting for each cluster. **(C)** Alluvial plot visualising the relationship of assigned cell clusters when defined based on RNA alone (left, 27 clusters), integrated RNA and ADT (center, 52 clusters), or ADT alone (right, 16 clusters).

Increased density of tumour infiltrating lymphocytes (TILs) is generally associated with improved prognosis, and the success of immunotherapies is often attributed to the modulation of TIL activity towards an anti-tumour response (Bruni et al., 2020). Previous studies of the breast cancer TME have identified important T cell and innate lymphoid cell (ILC) populations using fluorescence-based protein assays that have yet to be identified in scRNA-seq studies (Azizi et al., 2018; Janssen et al., 2020; Ruffell et al., 2012; P. Savas et al., 2018; Wagner et al., 2019). We identified several CD3+ cell clusters in which protein data played a particularly large role in clustering (elevated protein modality weighting; Figure 1B - asterisk marked) and which have a low silhouette score, indicative of poor cluster distinction and stability when calculated using RNA alone (Rousseeuw, 1987) (Figure S1F-G). The identification of such cell populations uniquely when using integrated analysis emphasises the utility of augmenting scRNAseq with proteomic measurements to enhance cellular phenotyping and delineate distinct cell populations. The elevated protein modality weighting in clustering of CD3+ cells suggest these cells would benefit the most from proteogenomic phenotyping, so we performed a targeted analysis of T cell and ILC populations.

### Targeted analysis of tumour-infiltrating lymphocytes identifies phenotypes which transcriptomics or proteomics alone cannot distinguish

A total of 7515 T/ILC cells passed QC, comprising 21 clusters, with each cluster including cells from at least 3 patients (Figure 2A-C; Figure S2A). Clusters were first stratified by protein expression of CD3 and TCRαβ, with high or mid-levels designated as conventional or /unconventional T cells and those with low levels classified as either natural killer (NK) cells or ILCs, depending on additional marker expression (**see Methods**; Figure 2D-E; Figure S2B). Two clusters with mixtures of both CD3 and TCRαβ high and low expressing cells were labelled as Lymphocytes (Lymph; C14 & C20). All CD3+ T cells were segregated based on CD8 or CD4 expression and assessed for expression of unconventional T cell markers or “NK-like” markers (see Methods; Figure 2D-E; Figure S2B). Top differentially expressed RNA and protein features for each cluster are shown in Figure 2E and **Table S3**. Each cluster was also scored for activity level (quiescent, low, mid or high) using a combination of published gene signatures (Szabo, Levitin, et al., 2019), total ADT abundance, RNA abundance, ribosomal content (Wolf et al., 2020), and known markers of T cell activation such as CD69, IFNG, GZMB, ZNF683/Hobit, PD-1, CD45RO (Cano-Gamez et al., 2020) (Figure 2B-C). Lastly, we collated gene signatures of commonly described T cell states, sourced from previously published studies (**Table S4**), with the aim to associate the observed clusters with an established lymphocyte effector function (Figure S2J).

**Figure 2:**
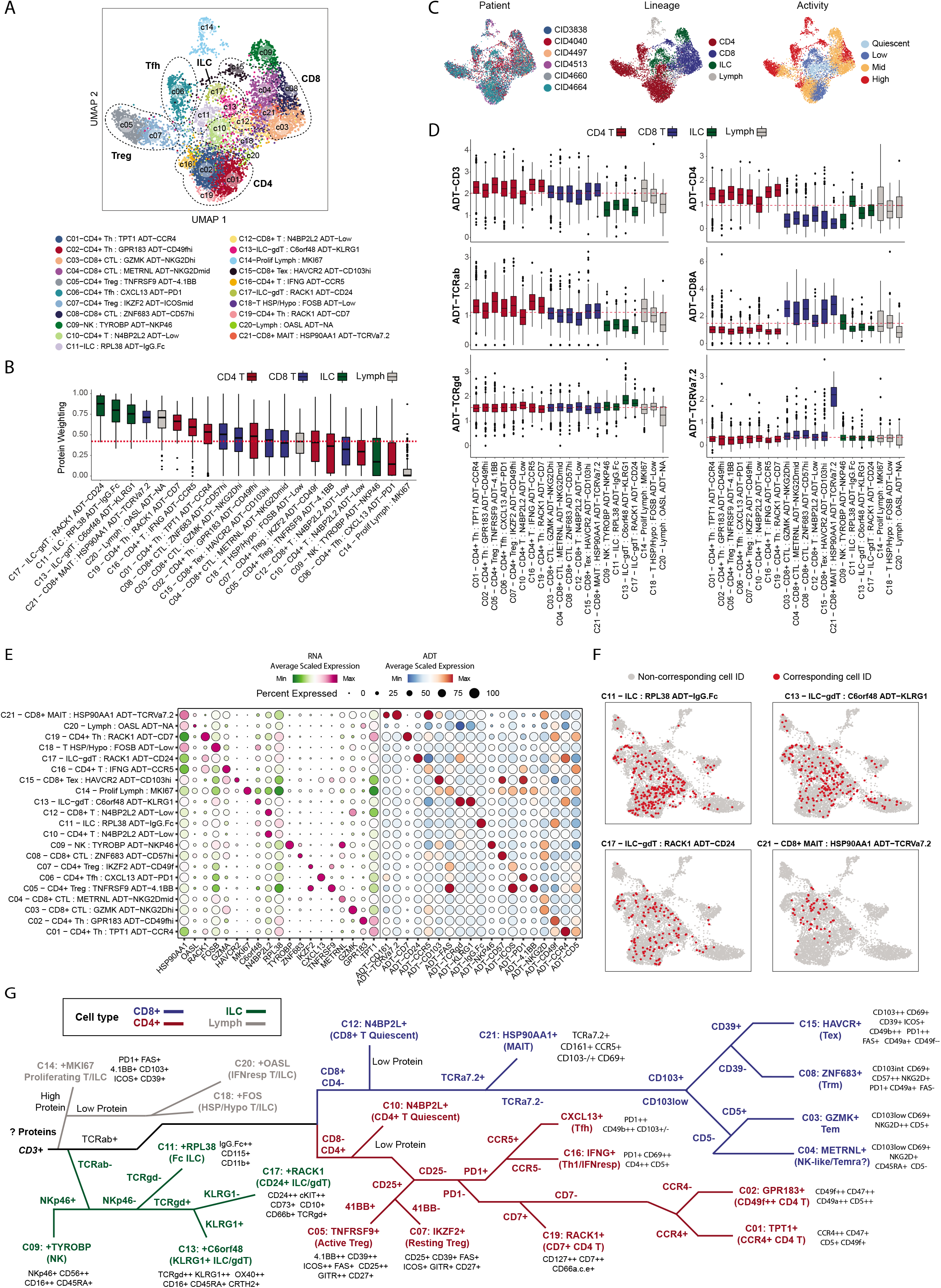
Targeted proteogenomic analysis of T and innate lymphocyte cells improves characterisation of TILs and identifies cell types which cannot be identified by RNA or protein modalities alone. **(A)** UMAP visualisation of integrated clustering of T and innate lymphoid cells derived from 6 human breast cancer samples. Clusters are numbered in order of decreasing cell number. **(B)** Protein weighting score, indicating the weighted proportion of ADT-derived features which contributed to the nearest neighbour calculation for the partitioning of cells into clusters. Increasing scores indicate increasing ‘weight’ of protein markers in the definition of a given cluster. Dashed red line indicates the median protein weighting score across clusters (0.451). Boxplots middle line marks the median value, the lower and upper hinges mark the 25% and 75% quantile, the whiskers correspond to the 1.5 times the interquartile range, the black dot’s mark outliers. **(C)** UMAP plots of cells in (A) colored by the following from left to right: patient sample, broad cell lineage and transcriptional activity score (see Methods). **(D)** Expression of lineage-defining ADT markers across T/ILC clusters, colored by assigned lineage and ordered by cluster size for each respective lineage subset. Boxplots middle line marks the median value, the lower and upper hinges mark the 25% and 75% quantile, the whiskers correspond to the 1.5 times the interquartile range, the black dot’s mark outliers. **(E)** Dotplot of expression (cluster average) of the RNA and ADT markers used to annotate each cluster. **(F)** WNN-derived annotations of clusters C11-ILC, C13-ILC/gdT, C17-ILC/gdT, and C21-CD8+ MAIT cells (red) projected onto the UMAP generated solely by RNA data. **(G)** Protein selection decision tree towards profiling of identified gene expression profile. ‘Low protein’ marks clusters which have have low enrichment of ADT-derived UMIs, ‘High protein’ marks cluster which have a high number of ADT-derived UMIs enriched.

Integrated analysis generated 7 additional lymphocyte clusters compared to analysis of RNA expression alone (Figure S2D-E). Three of these 7 clusters consist of CD3^low^ TCRαβ^low^ ILCs with a resting transcriptional profile, distinguishable by their expression of innate cell protein markers such as KLRG1, OX-40, cKIT, CD112 (PVR), IgG.Fc, or gamma-delta T cell receptor pairs (TCRgd), which are designated as C11-ILC:*RPL37* ADT-IgG.Fc, C13-ILC-gdT:*C6orf48* ADT-KLRG1 and C17-ILC-gdT:*RACK1* ADT-CD24 (Figure 2D-E; Figure S2B). These clusters are likely to reflect a genuine cell state as they show no abnormal gene contaminants from cells of other lineages, are low in mitochondrial genes, have nominal housekeeping gene expression, and show a negligible level of mouse transcript (ambient control) or isotype control ADTs (Figure S2H-I). We also identify an unconventional T cell cluster that is CD161^high^ TCRVa7.2^high^; canonical markers of mucosal associated invariant T cells (MAIT) which have previously been implicated in anti-tumour immunity (Petley et al., 2021), which we designated as C21-MAIT:*HSP90AA1* ADT-TCRva7.2 (Figure 2E). These 4 clusters have a low RNA-derived silhouette score and a high cluster similarity score, both of which reflect difficulty in identifying these phenotypes from transcriptomics alone (Figure S2F-G). Indeed, when these cell annotations were plotted onto UMAP space generated solely from RNA data, they were found to be dispersed, further supporting that transcriptomic data alone is insufficient for their demarcation (Figure 2F).

Multi-omic integration also significantly enhanced our ability to phenotype CD4+ T cells and cell states characterised by low transcriptional activity (Figure 2C; Figure S2E-G; **Figure SJ**). For instance, we were able to stratify CD4+ Treg:*FOXP3* cells into transcriptionally active (TNFRSF9^high^ ADT-4.1BB^high^) and resting (IKZF2^high^ ADT-ICOS^mid^) clusters, which were found to exhibit a gene expression profile similar to “Suppressive” and “Resting” Tregs respectively (Guo et al., 2018)(Figure S2E), previously characterised using Smart-seq2 which provides greater gene depth per cell compared to 10X Chromium (Wang et al., 2021). Our analysis also delineated CD4+ Th cells into 4 clusters of differing activation status, differentiated by their expression of T cell effector activation markers ADT-CD45RO and ADT-CD28, naïve markers such as CCR7 and IL7R and T cell resting enrichment scores (Figure S2E; Figure S2B-C). These were labelled, in order of low to high transcriptional activity: C02-CD4+Th:*GPR183* ADT-CD49fhi, C01-CD4+Th:*TPT1* ADT-CCR4, C19-CD4+Th:*RACK1* ADT-CD7, C16-CD4+T:*GZMA* ADT-CCR5. Integrated clustering also enabled us to separate a transcriptionally quiescent T cell cluster weakly positive for CD4 and CD8 transcripts (C04-T:*N4BP2L2* ADT-LE) into distinct CD4+ (C10-CD4+T:*N4BP2L2* ADT-LE) and CD8+ (C12-CD8+T:*N4BP2L2* ADT-LE) clusters (Figure S2E). While both of these CD8+ and CD4+ N4BPL2^high^ clusters lack any distinguishing protein markers outside of CD4 and CD8 expression (and are thus annotated as ADT-Low for “Low Expressing ADT”) (Figure 2E), they shared specific expression of long non-coding RNAs including *MALAT1*, *KIAA1551* and *N4BP2L2* and lack of expression of activation markers such as *CXCR4*, *CD69* and *NFKB1* (Figure S2B).

Proteogenomic analysis allowed us to associate protein markers with gene expression profiles previously described in scRNA-seq studies to play an essential role in TIL biology. To assist with visualisation of these markers we have provided a binary decision tree roadmap using the most distinctive protein markers of each cluster (6 for ILC, 7 for CD4+ T-cells and 6 for CD8+ T-cells), and an array of ADT markers enriched in each respective cluster (Figure 2G).

### Proteogenomic profiling refines markers of T cell activation, exhaustion and tissue residency

Experimental readouts typically provide a normative observation of feature expression; a feature is assigned as “high” or “low” in relationship to one another. Inconsistency can therefore arise when translating and/or integrating results across experimental assays or models where the features used to generate the comparative measurements are absent. We believe high-throughput cellular proteogenomics can provide a framework for a more standardised distribution of feature expression levels, particularly when performing cell type unbiased captures such as we have attempted with this dataset, and therefore improves standardisation of phenotyping. We first explored RNA and protein co-expression patterns of top differentiating cluster markers, including hallmark TIL features which previously have been described in literature to play an important role in defining their phenotypes (**See Methods**). As circulating T-cells infiltrate tissues they are reported to acquire the expression of either CD69 and/or CD103, markers sometimes used interchangeably to identify CD8+ tissue resident memory T cells (Trm) (Cibrian & Sanchez-Madrid, 2017; Lianne Kok et al., 2021; Okla et al., 2021; Szabo, Miron, et al., 2019). In the breast cancer TME, we find CD69 to be expressed across most T cell and ILC clusters (Figure 3A; Figure S3A), with increasing expression correlating with RNA signatures of TIL activation (Figure 3B)(Cibrian & Sanchez-Madrid, 2017) and with CD48 and CD2, both markers previously found to be associated with lymphocyte activation outside the context of tumour biology (Figure 3A; **Table S5**)(Binder et al., 2020; McArdel et al., 2016). In contrast, CD103 was more restricted (but not exclusive) to CD8 T-cells cells, proliferating cells and NK cells (Figure 3A; Figure S3A). Furthermore, we find that CD103 only modestly correlates with CD69 expression (Figure 3A; r=0.29, p = <0.001) (**Table S5**), with the exception of NK cells (r=0.58, p = <0.001) and exhausted CD8 T-cells (r=0.59, p = <0.001). We further observed CD103 can inversely correlate with CD69 in certain CD4+ T-cell states such as with Th1-like CD4 IFNG+ expressing cells (C16 - CD4+ T : *IFNG* ADT-CCR5 - r=-0.57, p = <0.001). These data suggest that in human breast cancer TILs, CD69 and CD103 cannot be used interchangeably and may be markers of more complex T cell phenotypes or states. Instead, we found the expression of marker 2B4 to correlate most frequently with CD103 across all TILs (r=0.48 p = <0.001) (Figure 3A)(**Table S5**).

**Figure 3:**
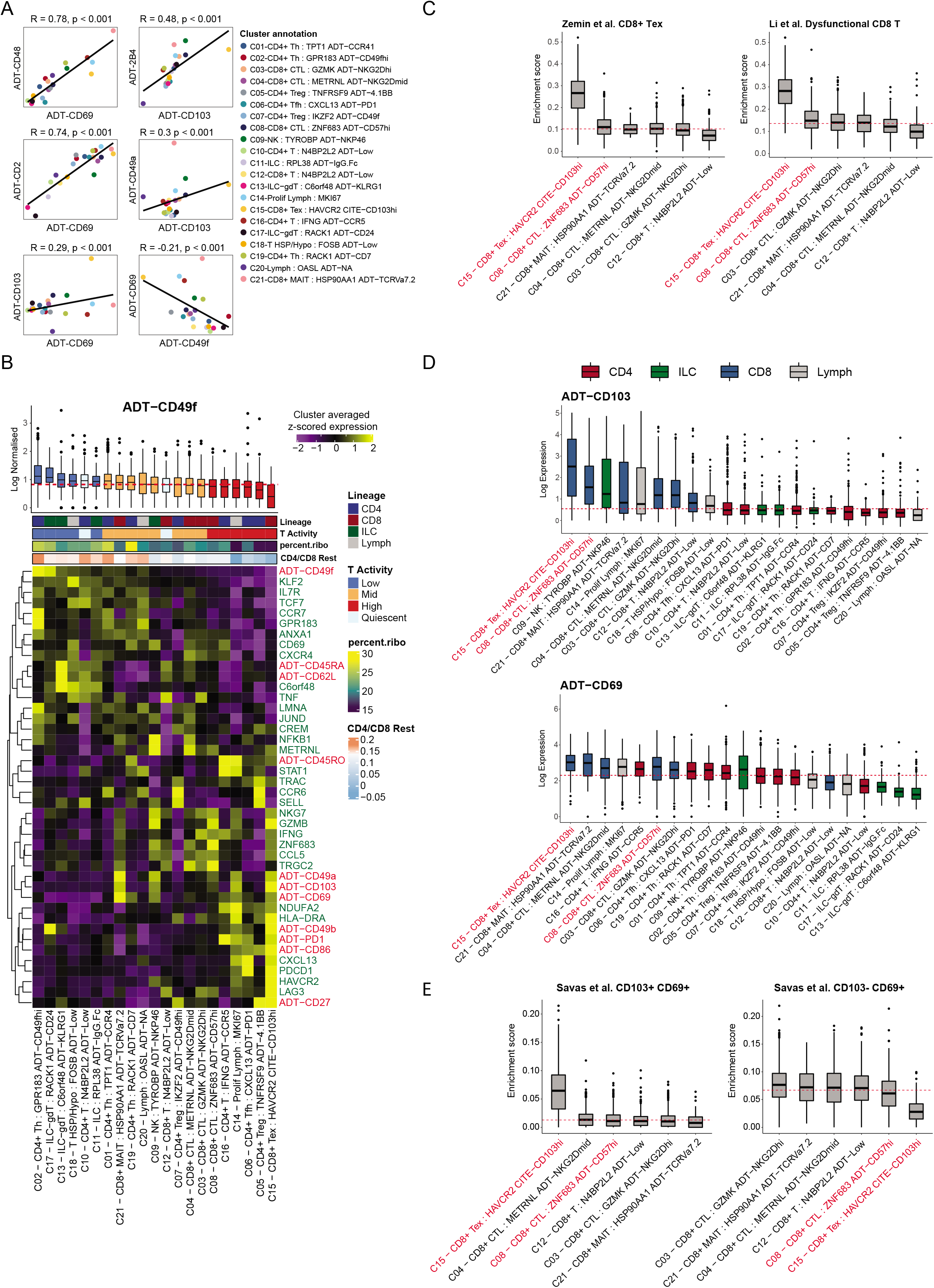
The expression of protein and RNA features of TIL activation and tissue residency markers in breast cancers. **(A)** Pearson correlation coefficient values of select pairing of ADT markers stratified by cell cluster. The global Pearson coefficient value for each ADT pair is provided, please see **Table S5** for Pearson coefficient values for each cluster. **(B)** RNA & ADT co-expression heatmap of features (rows) known to be relevant to activation, migration, exhaustion and tissue residency of T cells. Red text highlights ADT markers and green text highlights RNA markers. Boxplots at the top of the heatmap show the protein expression of CD49f grouped by T cell activity (see Methods), sorted from high to low (left to right). Boxplots middle line marks the median value, the lower and upper hinges mark the 25% and 75% quantile, the whiskers correspond to the 1.5 times the interquartile range, the black dot’s mark outliers. Percent_ribo refers to the proportion of ribosomal counts relative to all other expressed genes. “CD4/CD8 rest” is a published signature score based on (Szabo, Levitin, et al., 2019) that reflects enrichment of genes associated with inactive/resting CD4 & CD8 T cells. **(C)** AUCell enrichment score of signatures derived from Zemin et al. (Zheng et al., 2021) pan cancer CD8+ T-cell exhausted (Tex) and Li et al. (Li et al., 2019) dysfunctional CD8 T-cell, sorted from high to low (left to right). The top 50 differentially expressed genes were used towards calculating enrichment score. Red line indicates the median signature score value across all clusters. Red text indicates populations discussed in the text. Boxplots middle line marks the median value, the lower and upper hinges mark the 25% and 75% quantile, the whiskers correspond to the 1.5 times the interquartile range, the black dot’s mark outliers. **(D)** Log normalised expression of ADT markers CD103 and CD69 across T-cell and ILC clusters, sorted from high to low (left to right). Box plots are coloured by broad lymphocyte lineage. Red line indicates the median expression for each marker across clusters. Red text indicates populations discussed in the text. Boxplots middle line marks the median value, the lower and upper hinges mark the 25% and 75% quantile, the whiskers correspond to the 1.5 times the interquartile range, the black dot’s mark outliers. **(E)** Enrichment scores of gene signatures derived from Savas et al. study (Peter Savas et al., 2018) mapped into integrated CD8 T cell clusters. AUCell enrichment scores calculated from top 50 differentially expressed genes derived from bulk RNA-seq analysis of CD8 T-cells sorted by Flowcytometry into CD103+ CD69+ (left) or CD103-CD69+ (right). Boxplots are sorted from high to low (left to right) for each cluster. Red line indicates the median signature score value across all clusters. Red text indicates populations discussed in the text. Boxplots middle line marks the median value, the lower and upper hinges mark the 25% and 75% quantile, the whiskers correspond to the 1.5 times the interquartile range, the black dot’s mark outliers.

Increased TIL density can be used as a metric of tumour immunological status and is generally associated with good prognosis and response to immunotherapies (Maibach et al., 2020; Peter Savas et al., 2016). However, the presence of certain types of TILs, such as bystander TILs not specific for tumour antigens, resting or quiescent TILs, is likely to be less indicative of a strong anti-tumor response than infiltration by active tumour-specific CTLs (Scheper et al., 2019; Simoni et al., 2018). Therefore, it is important to find markers that can discriminate tumour-reactive TILs relevant to tumour immune status from inactive “passenger” T cells that can dilute the anti-tumor immune response. Unfortunately, lymphocytes with low transcriptional activity, such as resting T cells (clusters C04-T:*N4BP2L2* ADT-LE & C12-CD8+T:*N4BP2L2* ADT-LE) are challenging to phenotype with scRNA-seq (Figure 2; Figure S2F-G). This issue is exacerbated in tumours, where markers commonly used to identify antigen-naïve T-cells in blood such as CD45RA or lymph node homing markers CCR7 and CD62L (Martin & Badovinac, 2018; Payne et al., 2020) can be found on tumour-reactive T-cells and activated T-cells in ectopic lymphoid tissue(Ghorani et al., 2020; Wu et al., 2021; Zheng et al., 2021). Furthermore, some markers such as naive T cell marker CD44 are ubiquitously expressed at a high density across cell lineages and therefore challenging to use with cell-type unbiased CITE-Seq (Saturates CITE-Seq cDNA library) (Buus et al., 2021). Conversely markers commonly employed to assess T cell activation and/or migration state can be degraded by tumour dissociation (Autengruber et al., 2012). We therefore examined whether we could identify new protein markers that are enriched on resting lymphocytes in the TME. We found stem cell marker CD49f to be highly expressed on both CD4+ and CD8+ T cell clusters characterised by low transcriptional activity, particularly those with an elevated “resting” module score (**Methods**). Indeed, CD49f correlates more strongly with gene expressions markers of naive T cells (or early activated T-cells)(Gueguen et al., 2021), such as *IL7R*, *CCR7*, *TCF1* (encoded by TCF7) and transcription factor KLF2, which drives *S1PR1* and *CCR7* expression (L. Kok et al., 2021; Skon et al., 2013; Wolf et al., 2020) than protein levels of naïve T-cell markers CD62L or CD45RA (Figure 3B). We also found the proportion of ribosomal content to be a valuable indicator of TIL activity that can be inferred by scRNA-seq (high in resting cells), as previously suggested for PBMC-derived T cells (Figure 3B) (Wolf et al., 2020).

The measurement of both RNA and proteome has also provided us with a platform to directly evaluate phenotyping inconsistencies between transcriptomics and proteomics-based studies of breast cancer TILs, particularly relating to markers of tissue residency versus exhaustion. For example, a pan cancer T-cell analysis by Zeng et al. (Zheng et al., 2021) found naive CD8 T-cells to transition into exhausted CD8 T-cells (Tex) through two broad pathways, one which constituted granzyme K+ (GZMK+) expressing effector memory CD8 T-cells (Tem), and the other through tissue resident memory CD8 T-cells (Trm), which are governed by the residency-defining transcription factor Hobit (encoded by ZNF683+)(Park et al., 2019; Park & Mackay, 2021). In our dataset we identify two similar clusters of CD8+ T-cells, GZMK+ (C03 - GZMK ADT-NKG2Dhi) and ZNF683+ (C08 - ZNF683 ADT-CD57hi), prior to the acquisition of an exhaustion profile (C15 - HAVCR2 ADT-CD103hi) (Figure 3C; Figure S3B), which matches a phenotype described by others as dysfunctional (Figure 3C)(Li et al., 2019). Trm cells have been proposed by several studies across infectious disease and cancer models to be marked by CD103 and CD69 expression, including studies in human tissues of breast cancers (Peter Savas et al., 2018). However, integrated analysis revealed ZNF683+ Trm cells to be CD103intermediate. Instead, Tex cells (C15 - HAVCR2 ADT-CD103hi) had the highest expression of CD103 (2x higher than ZNF638+ T-cell cluster) (Figure 3D). Indeed, analysis of signatures derived from bulk RNA-Seq analysis of FACS-sorted human breast cancer TIL subsets reveals CD103+ CD69+ to primarily mark C15 - CD8+ Tex and not C08 - ZNF683+ Trm cells (Figure 3E). Instead, if we take high expression of ZNF638+ to mark tissue resident cells, we find CD8 Trm cells to be more precisely marked by low expression of protein markers CD39 or ICOS and elevated expression of either NKG2D and/or CD57 in addition to intermediate CD103 expression (Figure S3C; Figure 2E). Taken together, our analysis shows how the direct measurement of both RNA and protein modalities may resolve a discrepancy in the literature for the demarcation of Trm cells and Tex cells in the TME.

In summary, we find that integrated analysis of ADT and RNA co-expression is a valuable approach to nominate novel markers of the hallmark features associated with lymphocyte identity and activity in the tumour tissue context. Our dataset provides a rich resource to better associate cell surface protein expression with transcriptional patterns and cell states of TILs and can help clarify the functional consequences of established patterns of immune cell gene expression in the TME.

### Differentiated tissue resident T follicular helper cells can be classified by the expression of CD103 or *IGFL2*

Immune checkpoint inhibitors (ICI) targeting PD-1 have shown only a modest effect on survival in breast cancer patients compared to other cancer types such as melanoma, renal or lung cancer (Bruni et al., 2020; Wein et al., 2018). One of the primary therapeutic mechanisms of anti-PD-1 therapy is thought to be through prolonged activity of anti-tumour CD8+PD-1+ T cells, which would otherwise be inhibited by PD-L1/L2 expression within the TME (Wei et al., 2018). In a recent breast cancer single-cell atlas study, we found Tfh cells to be the most abundant PD-1-expressing cluster amongst all TILs across breast cancer samples (Wu et al., 2021), which remains consistent in this dataset (Figure 2E, C06−CD4+Tfh:CXCL13 ADT−PD1). The association of abundance of Tfh cells with improved tumour immunity have recently been described in several cancers, including breast cancer (Hollern et al., 2019b; Voabil et al., 2021; Zheng et al., 2021), partially attributed to their expression of the chemokine CXCL13 of which Tfh cells are a major source of (Wu et al., 2021; Y. Zhang et al., 2021). Tfh cells are known for their role in regulating the formation and activity of germinal centres, physiological microstructures found in lymphoid tissues that are essential for the development of high affinity antibody-producing B cells (Crotty, 2014; Ma & Phan, 2017). How Tfh cells behave within the TME remains under investigation. Tfh cells can acquire distinct phenotypes depending on the cancer (Zheng et al., 2021). Therefore, we next investigated whether enhanced phenotyping by integrated analysis of CITE-Seq can dissect breast cancer Tfh cells into clinically relevant subtypes.

After reclustering Tfh cells, we identified 3 clusters: CD4 Tfh:CXCR4, CD4 Tfh:CD103 and CD4 Tfh:IGFL2 (Figure S4A), all of which were enriched for expression of genes and pathways associated with Tfh cell features (Figure 4C; Figure S4B-D). Interestingly, we found a subset of these Tfh cells exhibited an exhaustion phenotype, reminiscent of dysfunctional/exhausted CD8+ T cells(Li et al., 2019; Wu et al., 2021), characterised by upregulation of CD103, 2B4, and CD49b proteins and *LAG3*, *CCL5*, and *HAVCR2* genes (Figure 4A-B; Figure S4E-F; Figure 3A). Surprisingly, GSEA of GO biological processes (n=6614 pathways, p < 0.05) shows CD103+ Tfh cells to correlate more strongly with CD8+ T exhausted cells than with other Tfh subsets (Figure 4D; Figure S4C). In contrast, a population of Tfh cells lacking CD103 expression were uniquely marked by elevated expression of the Neuromedin B (*NMB*) and Insulin growth factor ligand 2 (*IGFL2*) genes (Figure 4B; Figure S4E-F). *IGFL2* belongs to a family of 4 genes with unclear biological function that have been found to be infrequently and lowly expressed in various tissues, particularly skin, but upregulated during inflammation (Emtage et al., 2006; Lobito et al., 2011). We employed pseudotime analysis to explore whether breast cancer Tfh cells could be partitioned into stable transcriptional cell states along a trajectory of differentiation. Three stable states (Figure 4E-F) were identified, suggesting that activated Tfh cells (CXCR4^high^ BCL6^high^ IL7R^high^) can differentiate into either CD103+ Tfh (*HAVCR2*+ *LAG3*+), or *IGFL2+* Tfh cells (*NMB*^high^ *CXCL13*^high^) in the TME. When examining markers previously documented to be critical to the activity and differentiation of Tfh cell lineage, we find our segregated Tfh cell subsets have unique expression profiles, suggesting they each have distinct physiological roles within the TME (Figure 4F-G; Figure S4F). For example, the CD103+ Tfh cell subset acquires elevated expression of multiple chemokines including *CCL3*, *CCL4*, and *CCL5* known to be involved in the recruitment of inflammatory cells to the TME (Figure 4G)(Vilgelm & Richmond, 2019). This Tfh population also expresses higher levels of inflammatory mediators including IFNG, *IL17A*, *PRF1*, and *GZMB*, resembling Th1 or Th17 like Tfh which have been reported previously in different disease contexts including cancers (Morita et al., 2011; Singh et al., 2016; Zheng et al., 2021). In contrast, the IGFL2+ Tfh cell population showed specific upregulation of *IL-10*, *BTLA*, and *CXCL13*, markers important in the maintenance of GC reactions (Figure 4E; Figure S4F)(Cosgrove et al., 2020; Havenar-Daughton et al., 2016; Mintz et al., 2019; Xin et al., 2018).

**Figure 4:**
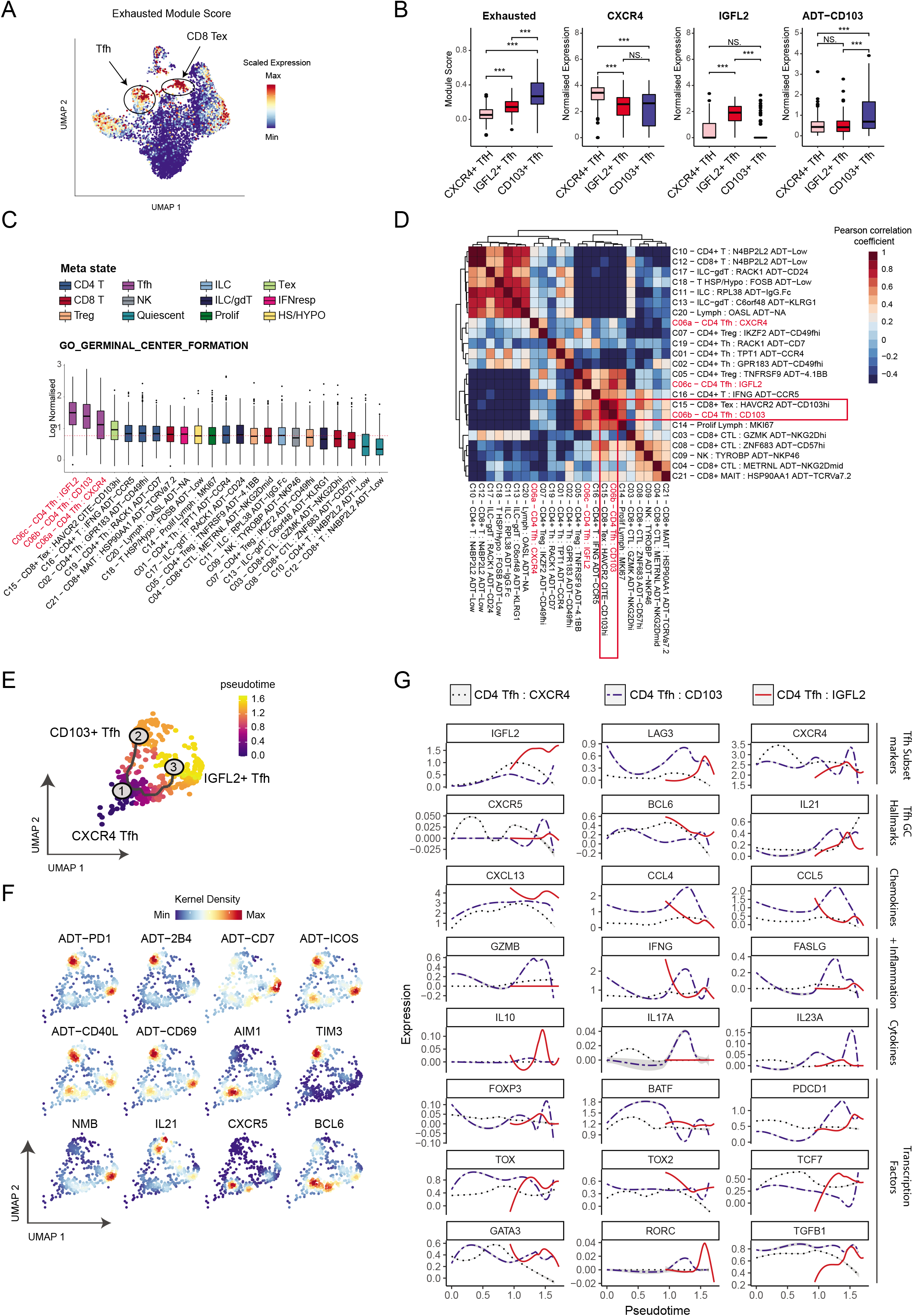
Integrated ADT and RNA clustering reveals novel subsets of Tfh cells in breast cancers. **(A)** Exhausted CD8+ Trm module score overlaid on the T cell/ILC WNN-derived UMAP (see Figure 2A). **(B)** Tfh cells divided into the 3 identified subclusters are plotted for, from left to right, exhaustion module score, normalised gene expression of *CXCR4* or *IGFL2*, and normalised protein expression of ADT-CD103. A two-sided t-test comparison between each cluster was performed, p-values are denoted by asterisks: *p < 0.05, **p < 0.01, ***p < 0.001 and ****p < 0.0001). Boxplots middle line marks the median value, the lower and upper hinges mark the 25% and 75% quantile, the whiskers correspond to the 1.5 times the interquartile range, the black dot’s mark outliers. **(C)** Gene set enrichment analysis boxplots for the GO biological process pathway “GO_GERMINAL_CENTRE_FORMATION” across T cells & ILCs, sorted from high to low (left to right). Red line indicates the median expression value across all clusters. Red text marks Tfh populations. Boxplots middle line marks the median value, the lower and upper hinges mark the 25% and 75% quantile, the whiskers correspond to the 1.5 times the interquartile range, the black dot’s mark outliers. **(D)** Pearson correlation heatmap of GO biological process pathways found to be significantly enriched (*n=6614, p < 0.05)* for each cluster. Red text marks Tfh populations. Red boxes highlight populations of interest outlined in text. **(E)** UMAP projections of pseudotime analysis using R package *Monocle3* (Cao et al., 2019) depicting the bifurcation of CXCR4+ Tfh cells (Node 1) into either the CD103+ exhausted-like state (Node 2) or an IGFL2+ state (Node 3). **(F)** RNA and ADT density kernel expression of known Tfh-relevant markers overlaid on UMAP projections derived from monocle3 pseudotime analysis (C) **(G)** Expression of known Tfh-relevant transcription factors, cytokines, chemokines and markers of subset differentiation within this study, stratified by subcluster, along the axis of pseudotime differentiation trajectory.

### Tissue CD103+ and IGFL2+ T follicular helper cells occupy distinct niches within the tumour microenvironment

Given the significant differences in gene expression phenotypes of Tfh cell populations, we hypothesised that these Tfh cell subsets may reside in unique tissue niches. We employed spatial transcriptomic data from 5 previously published breast cancer samples (2 ER+, 3 TNBC; Figure S5A) (Wu et al., 2021) to identify the location of Tfh subsets and co-location with other cell types. We observed that IGFL2+ Tfh cells are preferentially localised to regions characterised by high levels of immune infiltration, based on morphological examination, while CD103+ Tfh cells are preferentially localised to regions adjacent to cancer cells (Figure 5A-B). Assessment of co-localisation of these populations with other immune cell types shows that IGFL2+ Tfh cells often co-localise with both naive and memory B cells and dendritic cells, whereas CD103+ Tfh cells co-localise with memory B cells and macrophages (Figure 5C; Figure S5B). Higher resolution spatial analysis suggests that CD103+ Tfh cells are more frequently in proximity to proliferating T-cells and TREM2+ lipid-associated macrophages (Figure S5C-D), which express elevated protein levels of PD-L1 and are enriched in patients with poor disease outcome (Martina Molgora et al., 2020; Wu et al., 2021). In contrast, IGFL2+ Tfh cells are found to be significantly enriched proximal to CCR7+ CD4 T-cells, B cells and LAMP3+ DCs (mReg: DCs (Ginhoux et al., 2022)), antigen-elicited DC found to engage and regulate tumour reactive T-cells (Maier et al., 2020) (Figure S5C-D).

**Figure 5:**
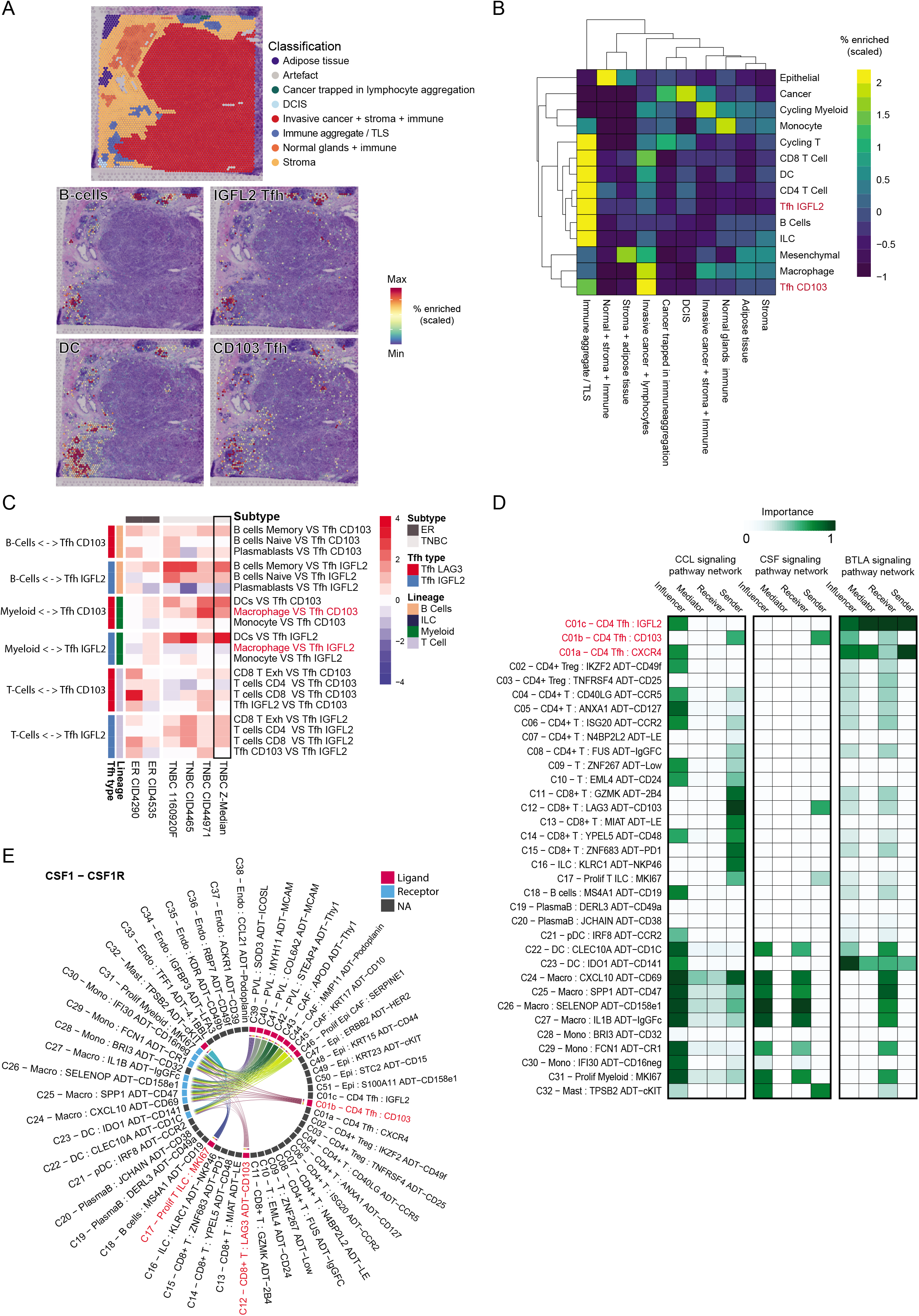
Localisation and signalling of tumour residing CD103+ and IGFL2+ Tfh subsets in breast cancers. **(A)** Enrichment scores for B cells, IGFL2 Tfh, CD103 Tfh and Dendritic cells (DC) overlaid on representative H&E images with pathological annotation shown for reference. **(B)** Heatmap of the enrichment (row scaled) of spatially deconvoluted cell types by pathological annotation as categorised by distinct morphological regions. The median value across 5 breast cancer samples from Wu et al. (Wu et al., 2021)(3 TNBC, 2 ER+) were used. **(C)** Pearson correlation heatmap of spatially deconvoluted cell pairs of interest. Red text marks populations of interest described in text. **(D)** CCL, CSF and BTLA signalling pathway network characterization across immune cell clusters. ‘Sender’ and ‘Receiver’ status reflects direct expression of ligands and receptors (agonistic or antagonistic). ‘Mediator’ and ‘Influencer’ quantifies the potential role in controlling receptor-ligand expression flow of the pathway within the system (here TME). Red text marks Tfh populations. **(E)** Chord Diagram representing the inferred cell-cell signalling of the CSF1-CSF1R pathway across all immune cells in the dataset. Red text marks populations of interest described in text.

To identify the signaling potential of Tfh cell subpopulations, we employed a receptor-ligand signalling pathway analysis. Using the expression of cognate receptors and ligands, signal events and directionality can be predicted using scRNA-seq data (Jin et al., 2021). A number of pathways were distinct between the two Tfh cell subsets (Figure 5D; Figure SD-F). Interestingly, the CD103+ Tfh cell subset had a unique CCL chemokine signalling profile reflecting elevated expression of CCL molecules and CSF1 (p < 0.01) (“sender” status, Figure 5E; Supp Figure 5F). They shared this profile with LAG3+ CD8+ T cells, which with proliferating T cells were the sole T cell secretors of CSF1 (Figure 5D-E; Figure S5H). The dual production of CCL and CSF1, critical ligands for macrophages, and the proximity to macrophages suggest a potentially unique role for this Tfh subset in interacting with macrophages within the TME. This role is further supported by enrichment of the GO biological processes’ “Macrophage activation involved in immune response” and “Chronic inflammatory response” in this subset (Figure S5E). In contrast, IGFL2+ cells have a significantly elevated engagement in BTLA signalling (Figure 5D, Figure S5F-G), a pathway active in restraining GC B cell selection and proliferation (Mintz et al., 2019).

### IGFL2+ Tfh cells are associated with poor prognosis in most cancers and enriched in anti-PD-1 poor responders in breast cancer

Previous studies associate the enrichment of Tfh cells with favourable prognosis across several cancers (Hollern et al., 2019a; P. Savas et al., 2018), and CXCL13-producing Tfh cells have been specifically implicated in survival and response to chemotherapy and immunotherapy (Ayse Bassez et al., 2021; Bindea et al., 2013; Gu-Trantien et al., 2013; Litchfield et al., 2021; Yuanyuan Zhang et al.). We therefore investigated whether one or more of the Tfh cell states identified in this study is specifically associated with clinical outcome. We generated a gene signature for each Tfh cell subset, based on the top differentially expressed genes in each population. Given their gene expression overlap with other T cells, we also included CD8+ Tex and Treg clusters in our analysis, in order to generate gene expression signatures unique to Tfh cell subsets (Figure S6A). Survival analysis shows the CD103+ Tfh cell expression signature to be significantly associated with improved survival in breast cancer (HR = 0.63, p = 0.0057) when compared to that of IGFL2+ Tfh cells (HR=0.85, p = 0.34), despite IGFL2+ Tfh cells found to express 2x more CXCL13.

As Tfh cells are one of the highest expressors of PD1, we reasoned that their activity may be influenced by therapies targeting PD1. We examined a scRNA-Seq dataset that explored the effect of anti-PD-1 checkpoint inhibitors on human breast cancers, in which patients were stratified by T cell clonal enrichment (expanders vs non-expanders) as a surrogate measure of anti-tumour activity levels (Ayse Bassez et al., 2021). We found that Tfh cells in this dataset could be similarly annotated based on either *IGFL2/NMB* (IGFL2+ Tfh) or *LAG3/HAVCR2* (CD103+ Tfh) expression (Figure S6B). Remarkably, the baseline abundance of both Tfh cell subsets was positively associated with response, with CD103+ Tfh cells showing a pronounced association with response, even greater than exhausted CD8 T cells, the presumptive target of anti-PD1 treatment (Figure 6B). When we explored the dynamics of Tfh subset abundance through treatment between non-expanders and expanders, we observed a significant increase of IGFL2+ Tfh abundance during anti-PD-1 treatment in patients who did not respond to treatment (NE; Figure 6C). Furthermore, we observed gene expression changes in IGFL2+ Tfh cells following anti-PD-1 treatment (Figure S6C), suggesting that anti-PD1 treatment may directly regulate their activity. Combined, these results indicate that the ratio of CD103+ Tfh to IGFL2+ Tfh cells is an indicator of ongoing tumour immunity, and that the ratio of Tfh cell subsets and their gene expression can be altered by anti-PD-1 checkpoint blockade. Therefore, although Tfh cells have been proposed as biomarkers of active anti-cancer immunity, we find a specific subpopulation of these cells associated with improved response to immunotherapy.

**Figure 6:**
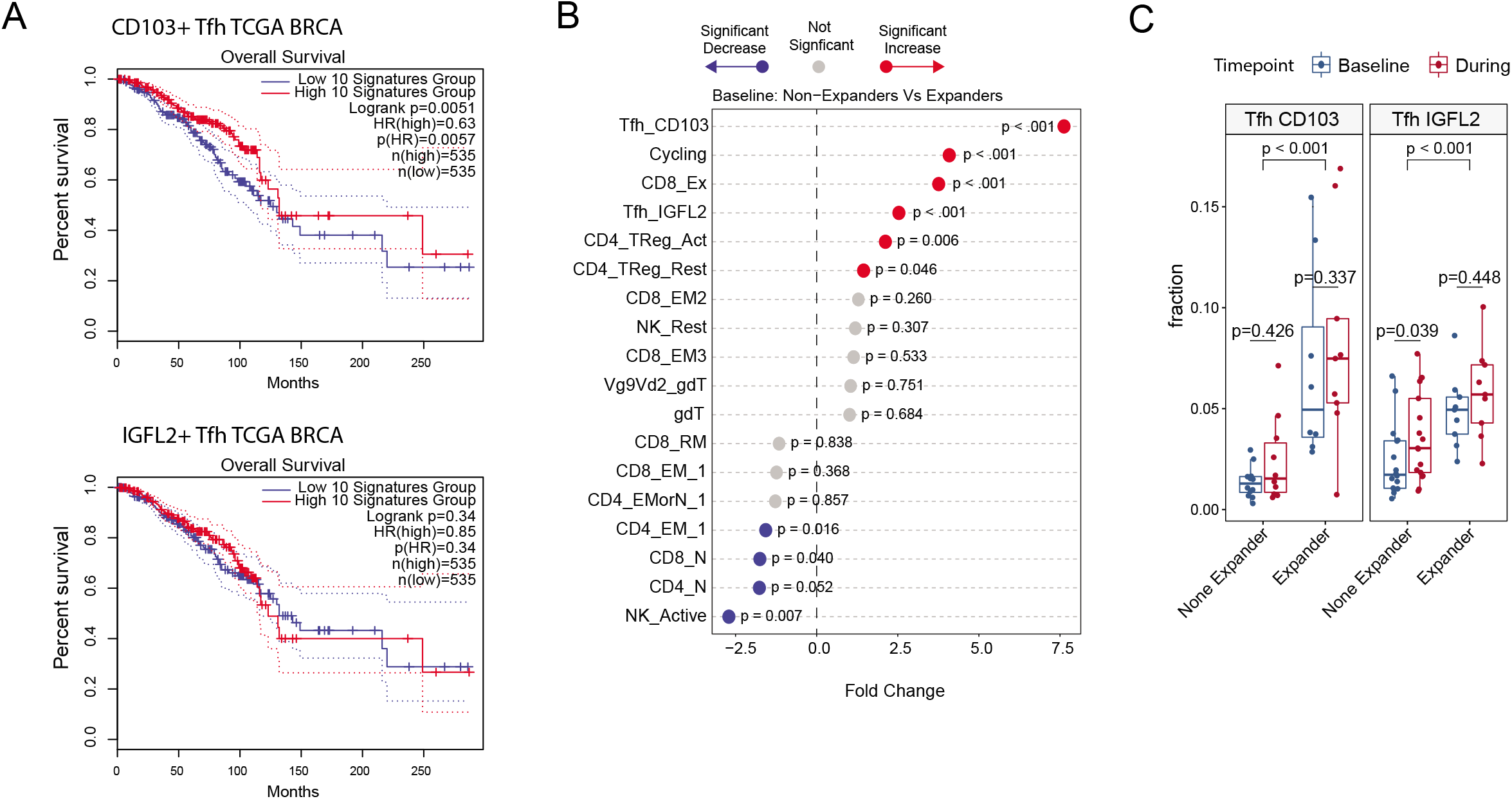
Tfh survival analysis and enrichment analysis of an anti-PD-1 breast cancer cohort. **(A)** Kaplan–Meier plots showing the associations between CD103+ Tfh and IGFL2+ Tfh signatures with survival in the TCGA cohort across all BRCA. The median group cutoff was used. P values were calculated using the log-rank test. The Hazard ratio was calculated as per cox proportional-hazards model. Dotted lines show 95% confidence interval. **(B)** Fold change in proportion of T cell subsets in pre-treated breast cancer patient samples classified as Non-expander (patients with low T clonal expansion) or Expander (patients with high T-cell clonal expansion), derived from Bassez et al. (2021), refined with Tfh subsetting. P-values calculated from non-parametric t-test (n=29) using package R package Speckle(Phipson et al., 2021). **(C)** Enrichment comparison of Bassez et al. (2021) breast cancer tumours sampled prior to (baseline) or during PD-1 treatment for none expanders (patients with low T clonal expansion) vs expanders (patients with high T-cell clonal expansion). Change of T-cell fraction of CD8 Tex, Tfh CD103 and Tfh IGFL2 subsets is shown. P-values are calculated from non-parametric t-test using R package Speckle (Phipson et al., 2021), see **Table S6** for n sampled for each condition or group. See Bassez et al. (2021) for further detail on the patient cohort (Ayse Bassez et al., 2021). Boxplots middle line marks the median value, the lower and upper hinges mark the 25% and 75% quantile, the whiskers correspond to the 1.5 times the interquartile range and dot’s mark outliers.

## Discussion

The incorporation of proteomic data with gene expression into a joint single-cell analytical framework for cell classification presents a promising avenue for improving the study of cellular behaviour within tissues, particularly cancer environments. Immunology has historically relied on a handful of cell surface protein measurements to delineate cell types; however, with the advent of single-cell RNA sequencing, it has become apparent that the array of functional cell types and states in the immune system is much more complex than previously appreciated(S. Cheng et al., 2021; Mulder et al., 2021; Triana et al., 2021; Zheng et al., 2021). Integrated proteogenomic resources like the one we report here allow for the precise linking of cellular transcriptomics to classical protein-based cell type dictionaries, such as those generated by the Immunological Genome Project (Aguilar et al., 2020). This approach thereby acts as a bridge to integrate a deep catalogue of classical immunological literature with cellular transcriptomic atlases.

Integrated RNA and ADT analysis was very effective in cellular phenotyping of the TME, allowing us to identify cell types and states which could not be defined using transcriptomics alone (Figures 1-2) (Azizi et al., 2018; Janssen et al., 2020; Ruffell et al., 2012; P. Savas et al., 2018; Wagner et al., 2019; Wu et al., 2021). The ability of transcriptomics alone to identify cellular subsets increases as a function of cell number, therefore integrated analysis provides substantial advantages in smaller studies using mid-throughput assays, for instance those using droplet partitioning which currently predominate in biomedical research. Integrated analysis particularly excelled in the phenotyping of cells with low transcriptional activity, such as ILCs and gamma-delta T cells, which lack defining gene expression modules but nonetheless express a discrete set of protein markers. The value of proteogenomic analysis in the identification of immune subsets appears to be in part a function of whether these cells are peripheral or TME-localized, particularly for cell types with cytotoxic properties. For instance, MAIT cells are easily identified among PBMCs and some cancer types using RNA data alone (Hao et al., 2021; Li et al., 2020). However in this study, and those generated by others in breast cancer (Azizi et al., 2018; Ayse Bassez et al., 2021; P. Savas et al., 2018; Wu et al., 2021; Yuanyuan Zhang et al., 2021), gene expression counts alone was insufficient to accurately distinguish MAIT cells from other CTLs (Figure 2F; **Figure SD-E**). Unconventional T cells and ILCs are increasingly shown to have an important role in regulating tumour immunity (Heinrich et al., 2022; Petley et al., 2021), improving methodology for their identification is therefore important. We believe integrated proteogenomics will be a major contributor in this regard.

TIL proteogenomics provided us with the tools to refine the protein markers commonly used for phenotyping using flow cytometry. We were able to explore new markers early along the trajectory of TIL activation, finding protein marker CD49f to be an alternative marker to CD62L or CCR7 for the identification of resting, naive-like or early activating tumour residing T-cells and ILCs. CD49f correlated more strongly than CD62L or CD45RA with early genes of TIL activation and differentiation such as *KLF2* (regulator of chemokine receptor expression and migration), *TCF7* (encodes TCF1), *IL7R*, and *CCR7* (Szabo, Levitin, et al., 2019; Wolf et al., 2020). CD49f (integrin a6) is a common stem cell marker, found on embryonic, mesenchymal, mammary, hematopoietic and cancer stem cells, and proposed to play an important role in self-renewal and differentiation of stem cells (Krebsbach & Villa-Diaz, 2017).

CD69 and CD103 are often used interchangeably to identify Trm cells despite both being imperfect markers for tissue residency (Lianne Kok et al., 2021). Previous studies have shown that Trm can lack the expression of CD69 and CD103, and their expression can also be found in circulating T-cells. We show that nearly all breast cancer TILs express some levels of CD69 (Figure S3A), with elevated expression as they increase transcriptional activity (Figure 3B). We also find CD69 to correlate poorly with CD103, in fact they were inversely correlated in some cell types (**Table S5**). Instead, we identified the protein markers CD2 and CD48 to be significantly co-expressed with CD69 (Figure 3A), both markers upregulated by activated circulating lymphocytes (Binder et al., 2020; McArdel et al., 2016). These data suggest that CD69 primarily marks TIL activation in human breast tissues. However, CD103 expression is largely restricted to CD8+ T-cells and NK, but significantly elevated (∼2X fold) by CD8+ Tex cells rather than *ZNF683+* T-cells which are most likely to be Trms (Figure 3D; (Caushi et al., 2021; Zheng et al., 2021). Instead, Trm were best identified by elevated expression of CD57 or NGK2D and an absence of CD39 or ICOS (Figure 2E; Figure S3C). This suggests that previous studies that relied on high CD103 expression for targeted analysis of Trm (Ganesan et al., 2017; L. Kok et al., 2021; Malik Brian et al., 2017; Nizard et al., 2017; Park et al., 2019; Peter Savas et al., 2018; Simoni et al., 2018) may have inadvertently included Tex cells, a population which we and others have found to have distinct expression of CD39 and resemble tumour-reactive CD8 T-cells (Duhen et al., 2018; Li et al., 2019).

Similar to FACS-based methods, a caveat regarding the application of this method is its reliance on individual markers for differentiating cell types. Additionally, performing CITE-Seq on dissociated tumour samples is technically challenging, and, as expected, we saw low signal-to-noise ratios for several protein markers, particularly those that are lowly expressed (Buus et al., 2020). This constraint may lead to inadequate differentiation of certain clusters in an unsupervised manner and may be particularly true for cell types where only one or two markers are used for differentiation, such as gdT cells (Payne et al., 2020). Therefore, identifying additional markers to delineate such cell types is crucial for further immune cell atlas efforts. We also believe the restricted number of cells typically captured by droplet-based platforms, the underrepresentation of certain cell types such as neutrophils and the lack of information about cellular granularity and size together suggest that this method cannot yet entirely replace FACS-based assays for phenotyping. Instead, we propose that ADT-derived cytometry complements FACS as a discovery tool for hypothesis generation, from which flow cytometric platforms can later be used to target cell types of interest for further analysis and quantitation. Our protein marker decision tree (Figure 2G) is a first step towards designing protein marker sets for FACS-based cell type identification.

Despite these limitations, CITE-Seq allowed us to generate new insights into the complex breast cancer immune TME. The canonical role for Tfh cells is to support B cells, yet we find a surprising abundance of Tfh cells in breast cancer relative to the proportion of B cells (B cells = 222, Tfh cells = 483) (Figure 1). As Tfh cells are the dominant PD-1 expressing cell type in the TME, and as PD-1 is the target of the most successful checkpoint blockade therapy currently in the clinic, we hypothesised that integrated analysis might reveal previously undescribed Tfh cell states with functions additional to their canonical role in germinal centre formation and operation. Indeed, we were able to identify three subsets of Tfh cells in the breast cancer TME - a baseline (CXCR4^high^/BCL6^high^/ IL7R^high^) state which differentiate into NMB^high^/IGFL2^high^ state and/or an exhausted-like HAVCR2^high^/LAG3^high^ state. Using CITE-Seq we confirmed the expression of CD103 and 2B4 by this latter population, markers typically associated with CD8+ Trm and Tex (P. Savas et al., 2018; Scott et al., 2019). Spatially-resolved transcriptomics showed distinct niches within the TME for the IGFL2+ and CD103+ Tfh cell populations. The IGFL2+ population was found predominantly in lymphoid aggregates resembling tertiary lymphoid structures and associating with B cells, suggesting a location and function similar to the physiological role for Tfh cells; conventional Tfh (cTfh). In contrast, CD103+ Tfh were found disseminated throughout the tumour, particularly in proximity to cancer cells and macrophages; cancer-associated Tfh (caTfh). The potential for unique interactions with distinct immune cell types by Tfh subsets was further supported by the expression of signalling molecules including CCL-family chemokines and CSF1 by CD103+ Tfh cells (Figure 5).

Immune cell phenotype is heavily influenced by the immediate cellular environment, with cancers in particular presenting diverse milieus in which TILs reside and receive signals, potentially leading to their acquisition of “neophenotypes” not found in the corresponding classical cell state or lineage. In our dataset, *LAG3*+/CD103+ CD4 Tfh cells and *LAG3+*/CD103+ exhausted CD8 T cells expressed many RNA signatures and protein markers in common, supporting the proposition that the TME can influence lymphocytes to acquire neophenotypes outside common lineage-associated phenotypes (Figure 5). Given their shared inflammatory gene expression, including CCL chemokines and CSF1, and co-localisation with macrophages, we hypothesise that both exhausted populations promote inflammation outside of lymphoid structures, which has recently been proposed in mouse models (Kersten et al., 2021). As both increased macrophage infiltration and increased exhaustion of T cells are strongly linked with patient prognosis (Cassetta et al., 2019; Foroutan et al., 2021), further studies on the downstream effect of exhaustion on adjacent immune cells within the TME is vital.

Survival analysis demonstrated the proportion of Tfh subsets to be of clinical importance, with CD103+ Tfh cell enrichment associated with improved survival, and improved response to checkpoint blockade in a breast cancer cohort (Figure 6). Conversely, the relative proportion of IGFL2+ Tfh was enriched in patients with low T cell clonal expansion following anti-PD1 treatment, a clinical feature associated with poor anti-tumour responses. The increased proportion of IGFL2+ cells, and changes in gene expression following treatment suggests that they are an important target of this immunotherapy. Recent reports have highlighted the importance of B cells and tertiary lymphoid structures (TLSs) in the response of diverse cancers to immunotherapy (Cabrita et al., 2020; Helmink et al., 2020). Our data provides the first evidence that Tfh subsets may make positive and negative contributions to the response to immunotherapy, warranting further investigation.

### Limitation of the Study

Generating good quality CITE-Seq data from dissociated tumours is challenging, noise is introduced either through non-specific binding of antibody oligos from lysed cells post staining or the through encapsulation of ambient antibody oligos with cells during oil droplet partitioning. This in addition to the affect that enzymatic dissociation can have on several epitope targets of antibodies, and the technical variation introduced by using a different antibody master mix for each capture. For the listed reasons, we believe the epitope data used in this study has a significant number of false negatives. Furthermore, as the sensitivity is reduced, we often opted to rely on cluster median or average expression when possible, rather than truly single-cell analysis.

Breast cancer is a heterogenous disease and can manifest a diversity of spatially organised structures. We explored only five datasets (3 TNBC and 2 ER+) for co-localisation of Tfh cells subsets. We believe a larger cohort is required to comprehensively understand their role in the TME, and in their manifestation depending on breast cancer subtype.

## Acknowledgements

We thank the following people for their assistance in the experimental part of this manuscript: The Garvan-Weizmann Centre for Cellular Genomics for their expertise in single-cell sequencing. Mr. Dominik Kaczorowski for his assistance in next-generation sequencing. This manuscript was edited by Life Science Editors.

## Author contributions

K.H. collected the tissue samples from surgery. G.A.-E and K.H prepared single-cell suspensions, G.A.-E performed CITE-Seq workflow. G.A.-E performed all analysis contained within this work except spatial deconvolution. A.A deconvoluted the spatial dataset. C.C. prepared and sequenced the ADT and RNA cDNA libraries. N.B., S.Z.W., D.R., T.W., J.R., E. M., provided technical assistance in interpretation or improvement of workflow. B. Y. provided TotalSeq antibodies. C.C.G, C. M., T. G. P., J. L., S.J., provided resources and biological interpretation of analysis and/or manuscript refinement. G.A.-E & A.S. conceived this work and wrote the manuscript.

## STAR Methods

### LEAD CONTACT

For any further information and requests should be directed to and fulfilled by the lead contact, Alexander Swarbrick.

### Data and Code Availability

The scRNA-seq data processed in this study is available to be explored and downloaded using the Broad Single-Cell portal at https://singlecell.broadinstitute.org/single_cell/study/SCP1793. All scripts used to process data and perform statistical analysis are available on https://github.com/Swarbricklab-code/BrCa_Integrated_proteogenomics. Raw FASTQ data can be accessed from the NCBI Gene Expression Omnibus database <u>GSE199219</u>. Any code used to visualise data is available from the corresponding authors upon reasonable request.

## Experimental model and subject details

### Patient material, ethics approval and consent for publication

The human breast cancer samples used in this study were collected following protocols x13-0133, x16-018, x17-155, x19-0496. Ethical approval for this study was acquired by the Sydney Local Health Districts Ethics committee, St Vincent’s hospital Ethics Committee, and Royal Prince Alfred Hospital zone. Consent for the use of samples in this study was obtained from all patients prior to collection of tissue, and data were de-identified as per approved protocol.

## Method Details

### Single-cell suspension generation of samples

Breast tissue was enzymatically and mechanically dissociated as per the Human Tumor Dissociation Kit (Miltenyi Biotec) protocol. The dissociated breast sample was then passed through a 100 µm MACS Smart Strainers (Miltenyi Biotec), topped up with RPMI 1640 10% FCS then centrifuged at 300 x g for 5 min. Supernatant was discarded, red blood cells were lysed using RBC lysing buffer (Becton Dickinson) for 5 minutes, then washed twice in PBS 10% FCS. All samples were cryopreserved in 10% DMSO, 40% RPMI 1640 and 50 % FBS solution then stored in liquid nitrogen until day of experiment, when samples were thawed in a 37°C liquid bath for 2 minutes, washed twice in RPMI 1640 10% FBS media, passed through a 100 µm strainer then resuspended in 100ul PBS 10% FCS media.

### Sample preparation and CITE-Seq antibody staining

TotalSeq-A antibodies (Biolegend, USA) compatible with 10X Chromium 3’ mRNA platform were used. The list of antibodies used for each sample are provided in the Supplement (**Table S1**). CITE-Seq was performed as previously described by Stoeckius et. al (Stoeckius et al., 2017) with the following modifications: Approximately 1 million cells per sample were resuspended in 95 ul of cell staining buffer (Biolegend, USA) with 5 ul of Fc receptor Block (TrueStain FcX, Biolegend, USA) for 15 min. Cells were then centrifuged at 350 x g for 5 min, supernatant discard, then 100ul of CITE-Seq mastermix (0.5ug of each Antibody) which was prepared earlier in that day with staining buffer (Biolegend, USA) was added to palleted samples. Cells were incubated for 30 min on ice, then washed three times. Approximately 3% 3T3 Mouse cells were then spiked into each sample as control, to estimate ambient RNA and ADT.

### Single-cell capture using 10x genomics chromium and sequencing

Cells for each sample were counted and confirmed to have > 80% viability using haemocytometer. Recovery of a total of 4000 to 6000 cells was the aim for each sample. Single-cell captures were performed using 10X Chromium Single-Cell 3’ v3 with exception to one breast tissue sample where Single-cell 3’ v2 kit was used (**Table S1**). Sample CID4676, CID4660, CID4664 were captured as one pool (multiplexed). Manufacturers protocol was followed in the preparation of RNA and ADT cDNA libraries. The cDNA libraries generated for each respective modality were sequenced separately on Illumina NextSeq 500. The following cycle settings were used for RNA cDNA libraries 28bp (Read 1), 91bp (Read 2) and 8bp (Index) and we aimed for 50,000 reads per cell. The following cycle settings were used for ADT cDNA libraries 28bp (Read 1), 24bp (Read 2) and 8bp (Index) and we aimed for 35,000 reads per cell.

### Single-cell RNA data processing

10x Genomics Cell Ranger (v3.0.4) was used to demultiplex BCL files to FASTQs, cell barcode demultiplexing, genome reference alignment (GRCh38 and mm10) towards generation of unique molecule identifier (UMI) count matrices. The CellRanger UMI counts from the “filtered barcode” list were used. All cells that have greater than 25% mitochondrial content and/or between 10% and 95% mouse mm10 aligned UMI’s were discarded as doublets, low quality cells, or as cells with increased ambient contamination. All mouse UMI counts except those expressed by the top 100 genes were removed prior to analysis. Samples CID4676, CID4660, CID4664 were demultiplexed using method “Souporcell” (Heaton et al., 2020) as instructed by developers and using default parameters. Genotype information for demultiplexing was generated by running UK Biobank Axiom array (ThermoFisher Scientific, Catalogue 902502) on each patient PBMC. The R package Seurat v4.0.4 was used for normalising, scaling, dimensionality reduction and cluster assignment using default parameters with two deviations, the first 40 principle components (PC) were used for dimensional reduction and to generate nearest neighbour graph, and an increased resolution of ‘1’ was used for clustering. Sample CID4676 was removed from all downstream analysis as it contained less than 50 cells.

### CITE-Seq data processing

The FASTQ demultiplexed reads for ADT libraries were assigned to each cell and antibody using package CITE-Seq count (v1.4.3, https://github.com/Hoohm/CITE-seq-Count), using the authors’ recommended parameters. Briefly, a cell barcode whitelist obtained from Cell Ranger “filtered” out for each sample was used for cell demultiplexing. A cell barcode levenshtein distance of 1 (--bc_collapsing_dist 1) and UMI distance of 2 (--umi_collapsing_dist 2) was allowed to be collapsed. The Antibody barcode list for TotalSeq-A (Biolegend, USA) used to demultiplex ADT is provided in **Table S1**. A levenshtein distance of 3 was permitted for ADT barcode demultiplexing (--max-error 3).

ADT counts were normalised using Seurat v4’s inbuilt centered-log ratio (CLR) transformation within cells (Margin 2). To determine which antibodies were enriched, we first constructed a nearest neighbour graph (‘FindNeigbours”), clustered cells on RNA (‘FindClusters’) at 1.2 resolution, the median absolute deviation of cells from each cluster was calculated and any cells that had 10-fold total ADT counts were discarded. To determine which antibodies are enriched within each sample, Seurat’s ‘FindAllMarkers’ across all clusters at 1.2 resolution after excluding mouse cells, was run for each individual sample. Only ADT features found to be enriched were used in the PC analysis, from which then the first 20 PCs were used towards dimensional reduction, graph construction and clustering analysis.

### Batch correction and Integration of RNA and CITE-Seq data

RNA and ADT assays were both first batch corrected across patients within each respective assay using Seurat v4 (4.0.4) prior to integration across assays. For RNA, the default parameters were used with the following deviation: the top 5000 anchor features were used in the step “FindIntegrationAnchors”, and the top 40 PC dimensions were used for the “IntegrateData” step. The top 40 PCs were similarly used for all subsequent steps; nearest neighbourhood calculation, cluster determination, UMAP calculation. ADT assay was processed similarly to RNA however only the top 20 PCs were used, and all ADT features (except Isotype controls) were used as anchors. The Batch corrected RNA and ADT matrices were then integrated using SeuratV4’s weighted-nearest neighbour (WNN), an approach which allows for simultaneous clustering of cells based on RNA and surface protein expression (ADT) (Hao et al., 2021). Integration was performed with developer recommended (default) parameters (*k = 20*) with the following modifications: The first 50 PC dimensions were used for RNA, and the first 20 PCs for ADT, in step “FindMultiModalNeighbors”. A resolution of 3.2 and algorithm 3 (SLM-smart local moving) was used for “FindClusters”. The majority of clusters were found to be present in most or all samples, with the exception of neoplastic epithelial cell clusters (C47-51) and clusters containing fewer than 50 cells, such as cluster C32-Mast:TPSB2 ADT-cKIT and C33-Endo:SEMA3G ADT-4.1BBL (Figure S1C).

### Silhouette score and cluster probability calculation

The silhouette coefficient was calculated using R package cluster 2.1.0. An euclidean distance matrix was generated from the first 40 PCs calculated from RNA assay alone, the same PCs which were used for dimensionality reduction and clustering as described above for processing RNA counts. The cell annotations were derived from the WNN approach, calculated from RNA and ADT assays as described above and shown for all cell lineages as in Figure 1 or targeted TIL analysis in Figure 2. To calculate the cluster probability, we used an approach previously described by Lun et al. ((Lun et al., 2016) Scran https://bioconductor.org/packages/devel/bioc/vignettes/scran/inst/doc/scran.html) which measures how many cells were partitioned into the same cluster after bootstrapping. We took the PCAs generated by Seurat’s v4 RNA assay analysis however with cell assignments to clusters set as per WNN of ADT and RNA modalities as described above. R Package Scran v1.18 was then used to bootstrap clusters (‘*bootstrapCluster*’) and to generate shared nearest neighbour graphs, and the paired co-assignment probability of cells to the same partition was evaluated using package igraph v1.2.6 function ‘cluster_walktrap’.

### Differential expression analysis, gene signature score modules, GO enrichment analysis

Differential gene expression analysis was performed using R package Seurat v4.0.4 function ‘FindAllMarkers’, using MAST v1.16.0 test (Finak et al., 2015). Module scoring for each gene signature was calculated using Seurat’s ‘AddModulScore’. The list of signatures used and their scource are available in **Table S4**. The cluster median module score of each signature was scaled 0-1 then visualised using spider plots using R package fmsb v0.7.0 (CRAN). R Package VISION v2.1 (DeTomaso et al., 2019) was employed to calculate enrichment of Gene ontology (GO) biological process’s, using immunological signature gene sets (c7.all.v7.2.symbols.gmt) derived from Molecular Signatures Database (MSigDB)(Subramanian et al., 2005). Either Complex Heatmap v2.7.11(Gu et al., 2016) or pheatmap v1.0.12 (CRAN) were used to visualise differentially expressed ADT, RNA, module scores and GO results. Seurat’s ‘DotPlot” function was used to visualise all dot plots. R package AUCell v1.12 (Aibar et al., 2017) was used to score CD8 T-cell Bulk RNA-seq signatures from Savas et al (P. Savas et al., 2018).

### T cell subsetting

Clusters were first stratified by their protein expression level of CD3 or TCRαβ, with high-mid levels designated as conventional/unconventional T-cells, and low levels classified as either Natural killer (NK) cells or ILCs, depending on expression of the markers ADT-NKP46, ADT-CD56, ADT-KLRG1, ADT-cKIT, ADT-TCRgd, GNLY and TRDC (Figure 2D-E; Figure S2B). Two clusters with mixtures of both innate and adaptive lineages were simply labelled Lymphocyte (Lymph). All CD3+ T cells were segregated based on CD8 or CD4 expression, assessed for expression of unconventional T cell markers such as ADT-CD161, ADT-TCRVa7.2 and ADT-TCRγδ, or markers that may signify “NK-like” features such as elevated ADT-NKG2D and ADT-CD57, TRDC, KLRD1, or GNLY (Figure 2D-E; Figure S2B). Top differentially expressed features for each cluster are shown in Figure 2E and **Table S3**. Each cluster was also scored for activity level (quiescent, low, mid or high) using previously described attributes of lymphocyte activation status such as published gene signatures (Szabo, Levitin, et al., 2019), total ADT abundance expressed by cell, RNA abundance and ribosomal content (Wolf et al., 2020), and known markers of T cell activation such as CD69, IFNG, GZMB, ZNF683/Hobbit, PD-1, CD45RO (Cano-Gamez et al., 2020) (Figure 2B-C). Lastly, we collated gene signatures of commonly described T cell states, sourced from previous published studies (**Table S4**), with the aim to associate the observed clusters with a previously ascribed lymphocyte effector function (Figure S2J).

### Spatial transcriptomic analysis

Visium patient sample counts and pathology notes were sourced from Wu et al. study (Wu et al., 2021). The single-cell dataset used towards deconvolution was similarly taken from Wu et al. study (Wu et al., 2021) however further processed to stratify Tfh cluster into CD103 Tfh and IGFL2 Tfh. To integrate the single-cell and spatial transcriptomics data (Visium), we used the software *stereoscope* v. 0.3.1 (Andersson et al., 2020). As input, the method takes raw UMI count data from the single cell and spatial transcriptomics experiments, together with cell type annotations for the former. From this, a single proportion matrix is produced. The matrix gives the proportion of each cell type (defined in the single cell data) at every spatial location. To improve the performance of stereoscope, we used a curated set of highly variable genes.

In order to reduce the runtime, we employed a subsampling strategy similar to that proposed in the original *stereoscope* manuscript. More specifically, we first defined a lower and upper bound (here, 25 and 250 respectively). Next, cells were sampled according to the following scheme: if a cell type had fewer members than the lower bound, we excluded it from the analysis; if a cell type had more or the same number of members as the lower bound, but fewer or the same number of members as the upper bound, we used all cells within the cell type; if a cell type had more members than the upper bound, we randomly sampled #[upper bound] cells from the cell type (without replacement).

*stereoscope* was run with the following parameter settings: batch size - 2048, number of epochs - 50000, These settings were used in both steps of *stereoscope*, i.e., the parameter inference step and the proportion estimation step. Default values were used for all other parameters.

Highly variable genes were extracted by applying a sequence of three functions from the scanpy suite (v. 1.7.2) to the single cell data. First, we normalized the gene expression data, then, the normalized values were log-transformed (using pseudocount 1), finally, the highly variable genes were identified from the transformed values. The exact function calls were:

scanpy.pp.normalize_per_cell(…,10e4)

scanpy.pp.log1p(…)

scanpy.pp.highly_variable_genes(…,n_top_genes=5000)

Where “…” represents an anndata object containing the relevant data.

### Trajectory analysis & Receptor-Ligand analysis

R package Monocle3 v 1.0 (Cao et al., 2019) was used to generate the pseudo-trajectory analysis involved with the characterisation of T follicular helper cells. Briefly, the RNA batch corrected and integrated matrix generated by Seurat v4 (4.0.4) were exported to build the CellDataSet object. Pseudotemporal analysis was then performed using default parameters as instructed by developers in their vignette.

The R package Slingshot v1.6.1 (Street et al., 2018) was used to generate the pseudo-trajectory analysis of CD8 T-cells. Default parameters were used, and UMAP input was derived from batch corrected RNA matrix values as generated by Seurat v4 (4.0.4). Trajectory overlay was mapped on cells clustered and annotated by Seurat v4 (4.0.4).

Ligand-Receptor analysis was performed using R package “CellChat” (Jin et al., 2021). Analysis was executed using default parameters as recommended by developers, as described by their vignettes.

### RNA and protein co-expression patterns of hallmark tumour infiltrating lymphocyte states and meta module analysis

Immune cell phenotype is heavily influenced by the immediate cellular environment, with cancers in particular presenting diverse milieus in which TILs reside and receive signals, potentially leading to their acquisition of “neophenotypes” not found in the corresponding classical cell state or lineage. As current immunotherapies primarily rely on cell surface expression of target proteins as biomarkers and therapeutic targets, it is important to better understand the TIL phenotypes that correlate with specific, often nonclassical, surface protein markers in the context of the TME. Thus, we explored the landscape of RNA and protein co-expression across our dataset in order to link hallmark gene expression features associated with functional lymphocyte commitment to ADT marker expression, independent of lineage. We used a semi-supervised data-driven approach to choose the most informative features to define T cell and ILC phenotype meta states; grouping cells by greatest variance (and similarities) into broad categories of TIL effector function. This included the top 5 differentially expressed genes plus all ADTs enriched in each cluster (MAST test; p_adj < 0.05, n = 178 genes; n = 63 ADTs), this included a curated panel of 30 differentially enriched (MAST test; p_adj <0.01) master transcription factors and/or genes previously associated with hallmark regulation of lymphocyte development, activation, and function in the TME such as TBX21(T-Bet), GATA3, EOMES, TOX, TCF1, TGFB1, BATF, and STAT3/5. Each feature value was scaled (Z-scored) for each respective assay then merged. An euclidean approach (sqrt(sum((x_i -y_i)^2) was applied to generate the dissimilarity matrix, which was then clustered using Ward.D2 using R package ComplexHeatmap v2.7.11. R Package igraph v1.2.6 was used to visualise the distances between each feature using a phylogenetic dendrogram. The partitioning of RNA and ADT features into clusters were generated using a centroid k-means approach. The appropriate number of selected k (number of clusters) was estimated by gap statistics using package fmsb v0.7.0. R package clustree v0.4.3 (https://doi.org/10.1093/gigascience/giy083) was used to visualise the impact of different *k* resolutions on cluster assignment. Clustering with this set of features revealed thirteen functional meta states defined by co-expressed genes and protein markers (Figure S7A-E). We observed trends consistent with the literature, such as the association between the expression of CD103 with GZMB and exhaustion gene HAVCR2, or between CD39 surface expression and genes upregulated by immunosuppressive T cells such as FOXP3+ Tregs (Supp Figure 7D-E). We also interrogated the co-clustering patterns of RNA and its cognate protein. We identify a few genes whose RNA and protein levels cluster closely irrespective of lineage, suggesting that either measurement is sufficient to capture robust expression information for these markers. Examples include PDCD1 and PD-1, IL2RA and CD25, CD4 and CD4 and CD8A and CD8A/B (Supp Figure 7D-E). However, the majority of genes tend to demonstrate poor correlation between RNA and protein. Importantly, numerous mRNAs are poorly detected despite high expression of their cognate protein, such as ADT-CD57 and ADT-CD7, as well as the group of ADTs that represent the primary cluster drivers of resting ILC cells, indicating that the assessment of such cell types with transcriptional methods alone will remain challenging.

### Survival analysis and patient cell type proportion analysis

Survival analysis of was performed using ‘GEPIA’, using the TCGA BRCA cohort http://gepia2.cancer-pku.cn/#index (Tang et al., 2017). The top 10 differentially expressed genes of each Tfh subset of interest were used and the median value was used as group cutoff. The hazards ratio was calculated as per cox proportional-hazards model and the p-value was calculated using log-rank test statistics.

R package speckle (0.0.2) (Phipson et al., 2021) was used to calculate statistical significance in the change of cell type proportion between patient groups or conditions. All Beassez et al. patients (A. Bassez et al., 2021) which were not assigned either as Expander (E) or Non-expander (NE) were removed from analysis (two patients’ pre-post samples were excluded). Change in composition was visualised by calculating the fraction of each cell type within each patient. A non-parametric t-test (n=29) was used to calculate the p-value of baseline change of composition between Expanders and Non-expanders. A non-parametric t-test was used to calculate p-value between each respective group comparison, *n* sample used for each comparison shown in Figure 6C can be found in **Table S6**. See Bassez et al. for further detail on patient cohort (A. Bassez et al., 2021).

### Statistics and reproducibility

No statistical method was used to predetermine sample size. The statistical significance for all differentially expressed genes or ADT were determined using MAST(Finak et al., 2015), and adjusted bonferroni corrected values were used. The Box plot centre line depicts the median value, the first and third line mark the 25% and 75% quantile, the whiskers correspond to 1.5x the interquartile range (IQR), and dots mark outliers. Details for any statistical tests performed are present in figure legends and in relevant method sections.

## KEY RESOURCES TABLE

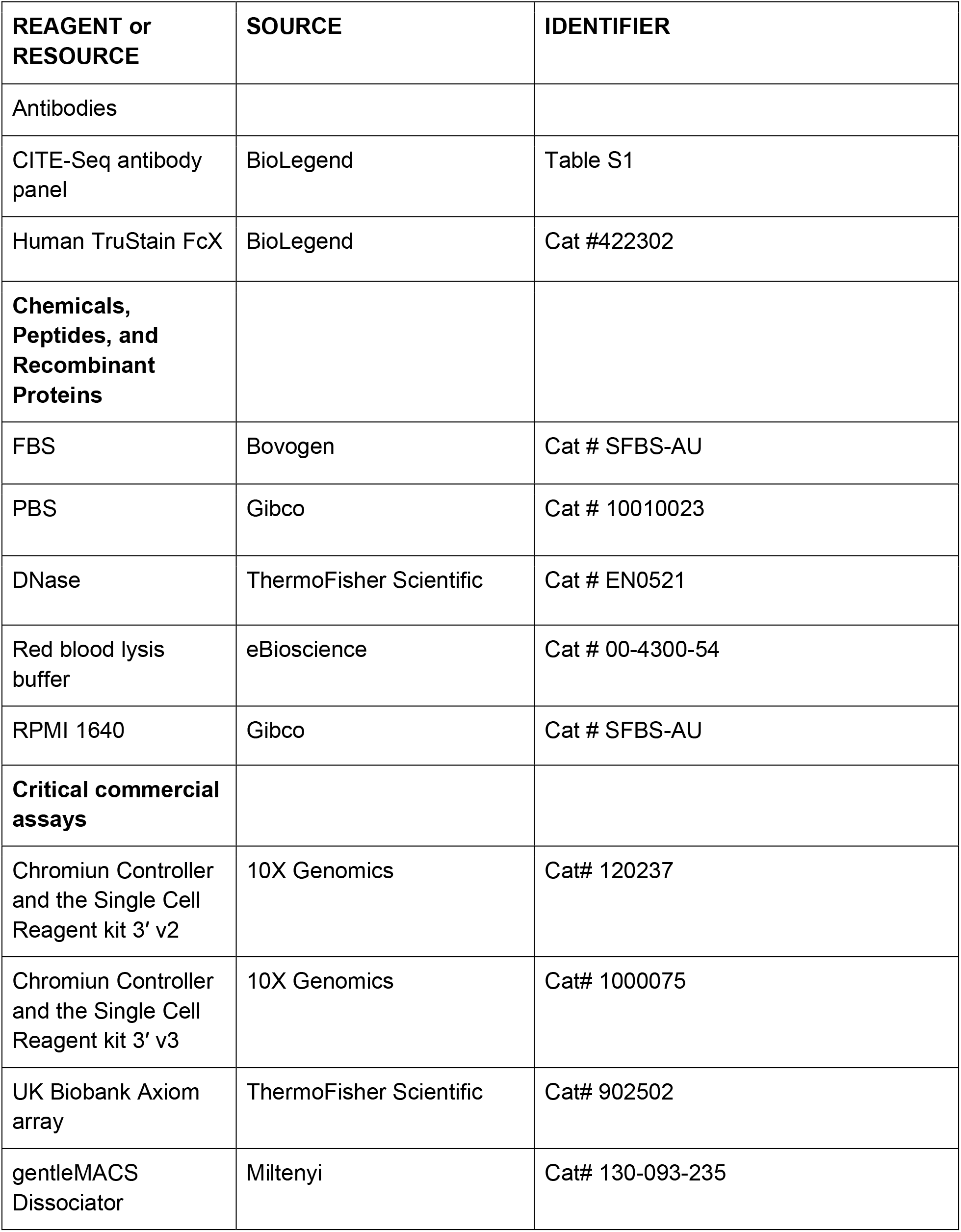
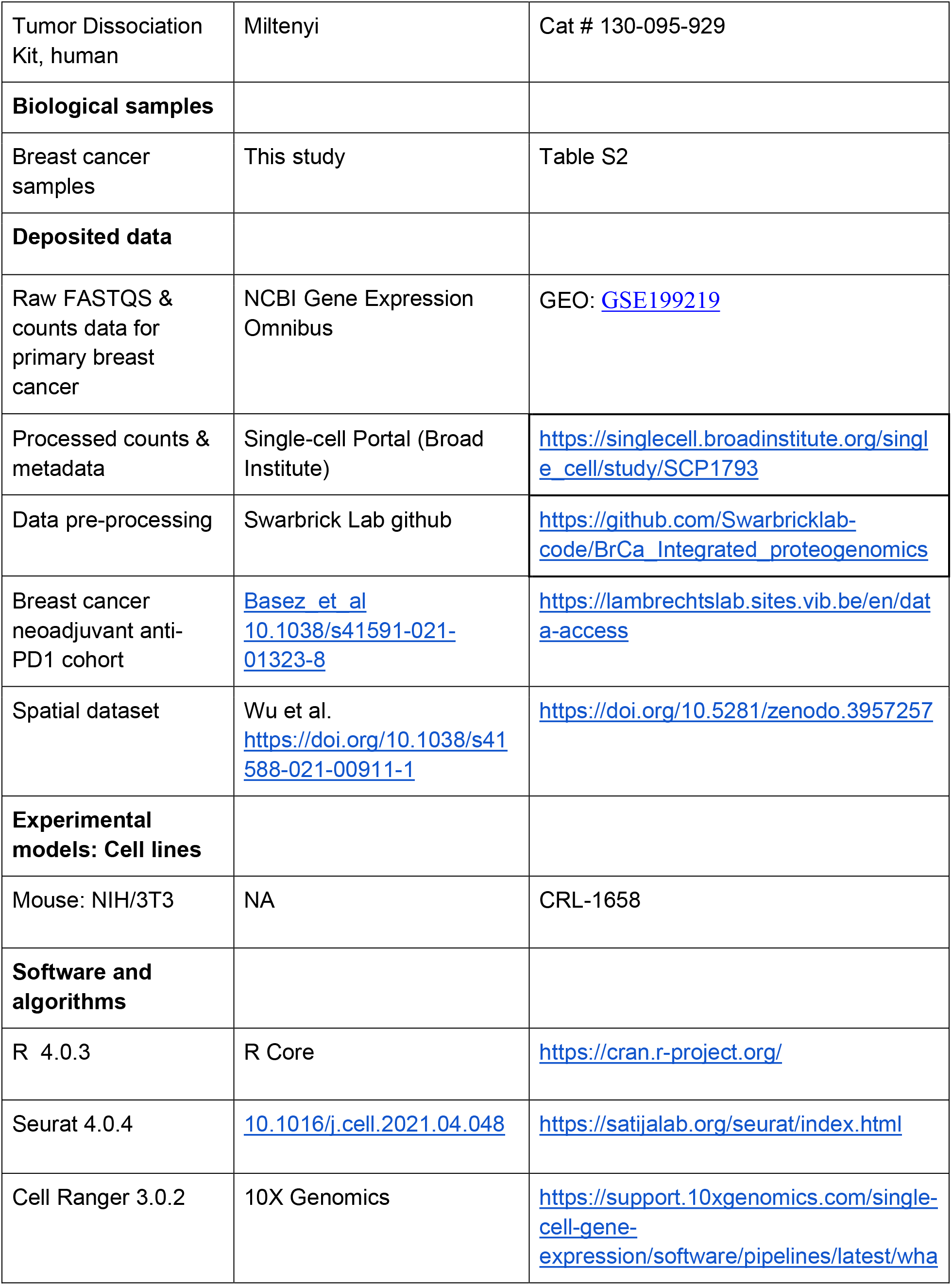
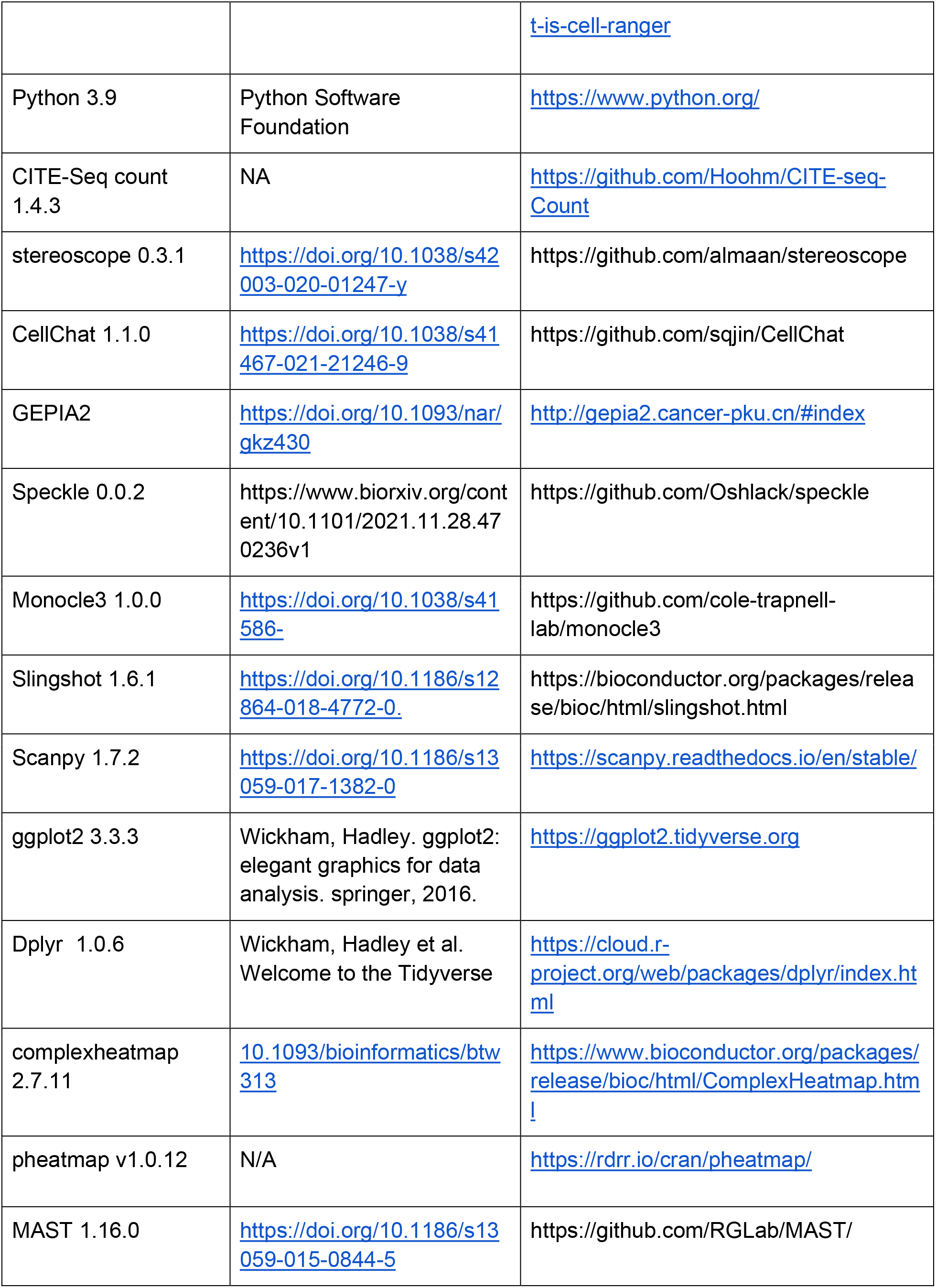
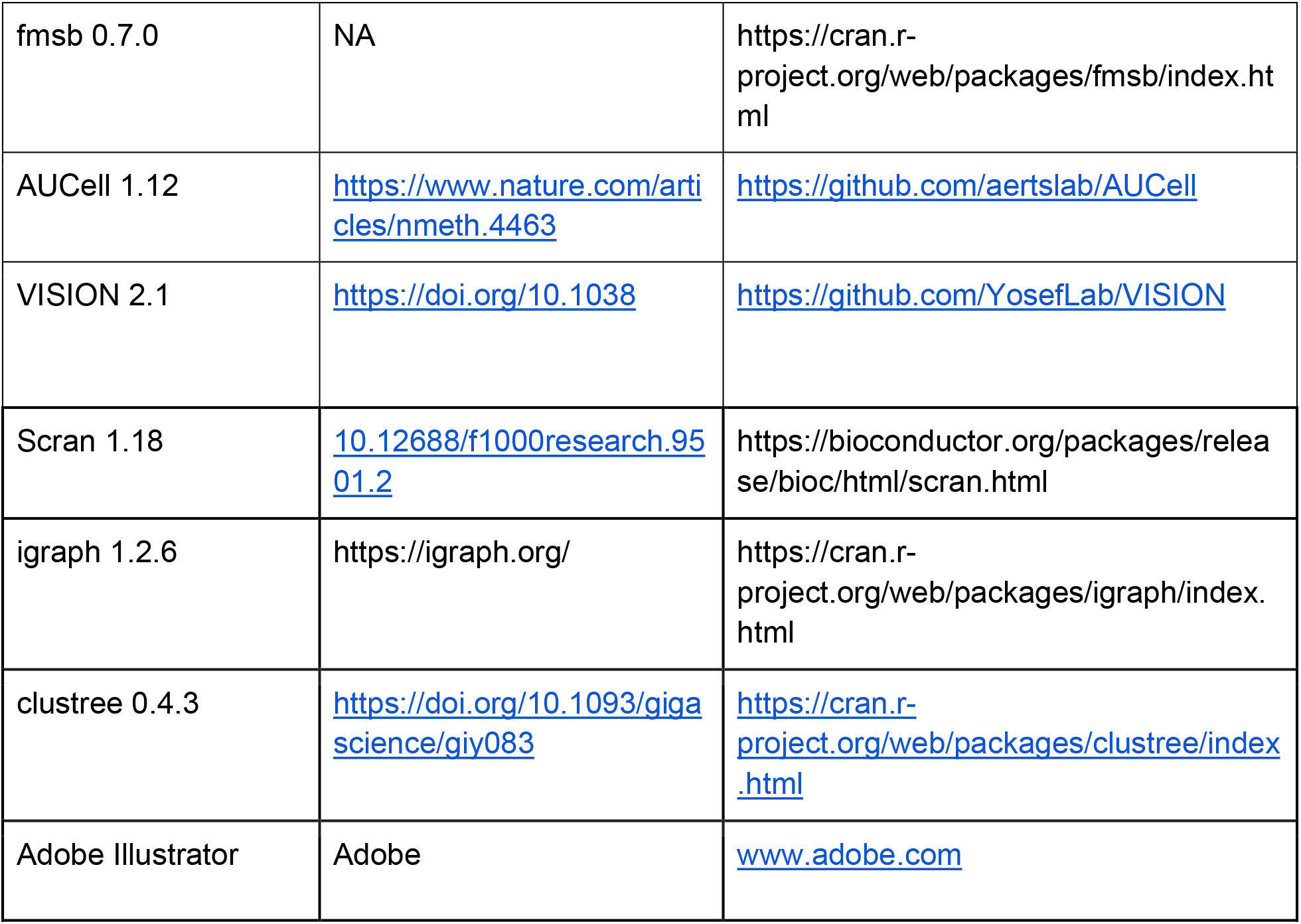

## Supplemental Information

**Supplementary Figure 1:**
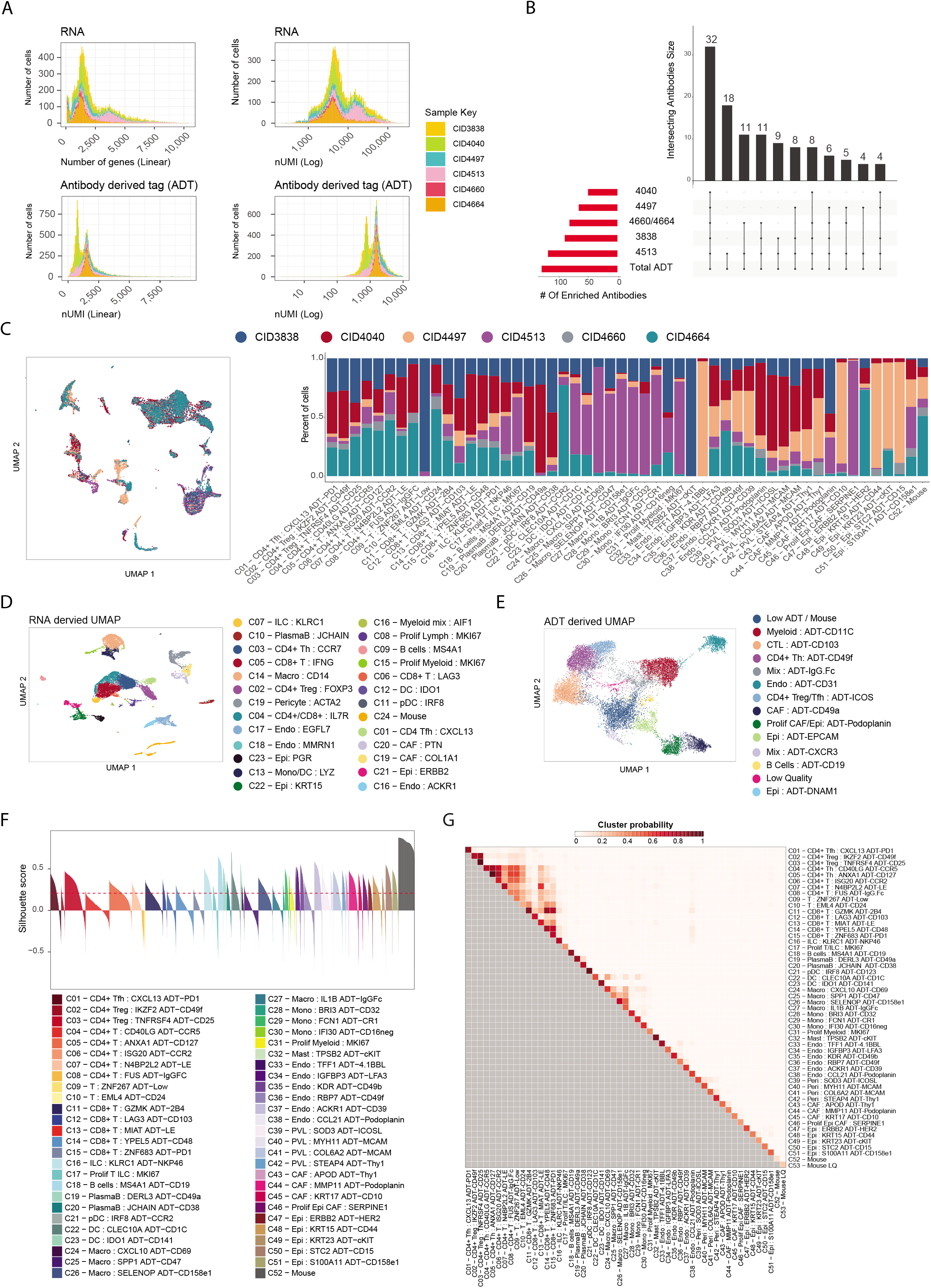
Quality control metrics and cluster information of CITE-seq data derived from 6 breast cancer samples. **(A)** Number of genes and UMI counts for both ADT and RNA assays, and mitochondrial content proportion for each sample, information of patient sample disease are available in Supplementary Table 2. **(B)** UpSet plot of CITE-seq antibodies found to be commonly enriched across samples. **(C)** UMAP of cells grouped by patient sample, and the proportion of each cluster derived from each patient. **(D)** UMAP of all cell clusters derived solely from RNA analysis. **(E)** UMAP of all cell clusters derived solely from ADT sequencing. **(F)** Silhouette score of each cluster for the evaluation of cluster stability. **(G)** Cluster stability matrix indicates the probability that a random cell from each paired cluster would be re-assigned to the same cluster following bootstrapping. High probability means paired clusters contain more related cells than not.

**Supplementary Figure 2:**
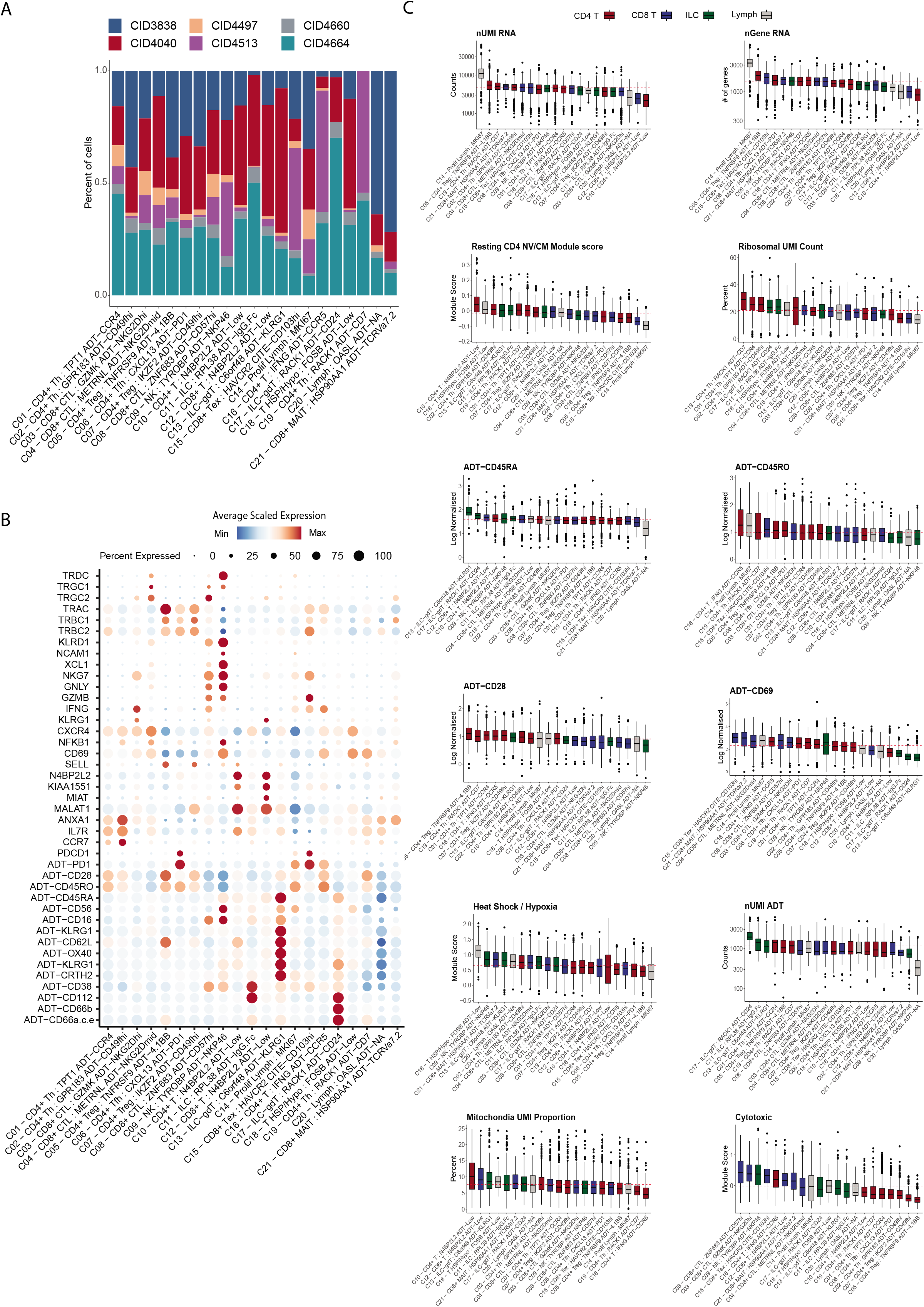
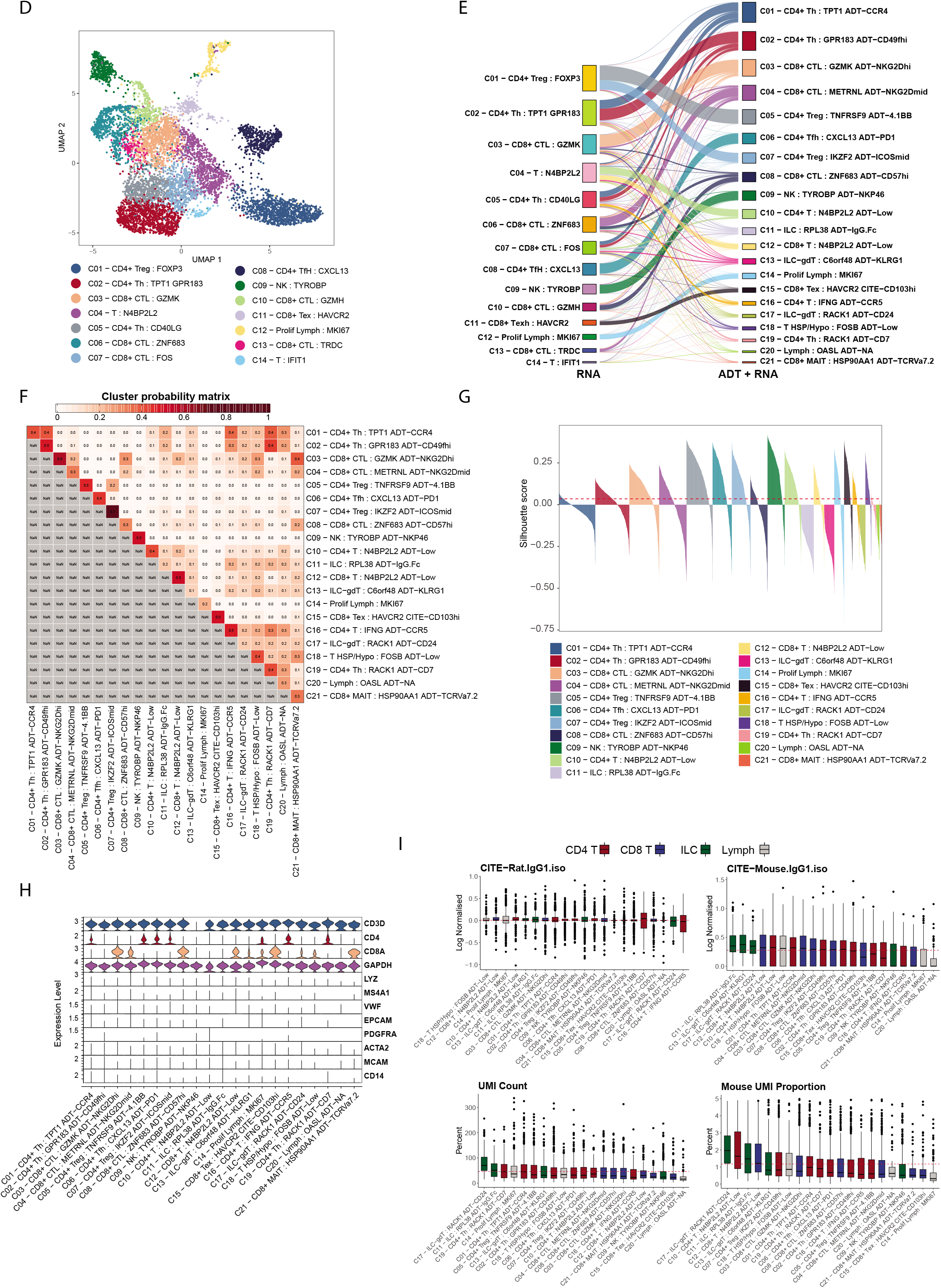
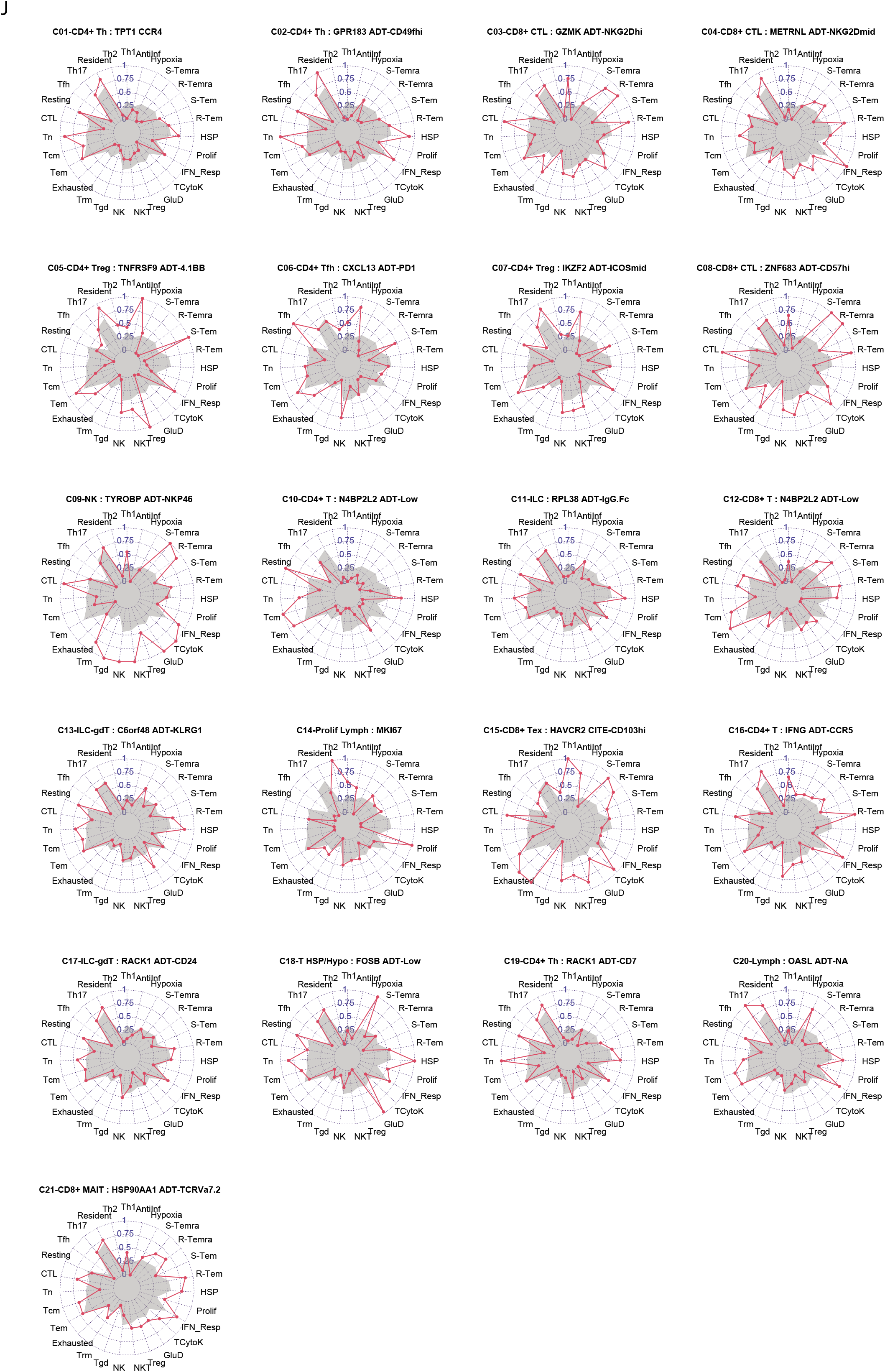
Supplementary information for RNA and protein Integrated clustering analysis across T cell and ILC populations. **(A)** Proportion of patient cells from each sample belonging to each T/ILC cluster. **(B)** Dotplot visualisation of the expression of RNA and ADT markers of interest on a z-score normalised scale. **(C)** Log normalised expression of factors associated with lymphocyte activity and naïve/memory activation status. Bar plots are coloured by lymphocyte lineage. nUMI refers to the number of unique molecular identifiers derived from the RNA assay, and nGene refers to the number of genes identified. Heat shock/Hypoxia and Cytoxic enrichment scores were calculated using Seurats “*ModuleScore”* using genes listed in Table S4. Red line indicates the median expression for each marker across clusters. **(D)** UMAP of the T cell/ILC dataset when clustered on RNA features alone. **(E)** Alluvial plot visualising the relationship of assigned cell clusters when derived from RNA alone versus RNA and ADT integration. **(F)** Cluster stability matrix shows the probability that a random cell from each paired cluster would be re-assigned to the same cluster following bootstrapping. High probability means paired clusters contain more related cells than not. **(G)** Silhouette score of each WNN-derived cluster calculated from RNA alone. Negative values indicate instability. **(H)** Violin plot showing normalised expression values of canonical markers of lymphocytes, myeloid, epithelial, and mesenchymal lineages. **(I)** Expression levels of isotype controls and mouse spike-in. Red line indicates the median expression. **(J)** Radar plot projecting the gene signature module scores, scaled 0-1, for each respective cluster. Grey silhouette marks the module score profile when averaged across all clusters combined.

**Supplementary Figure 3:**
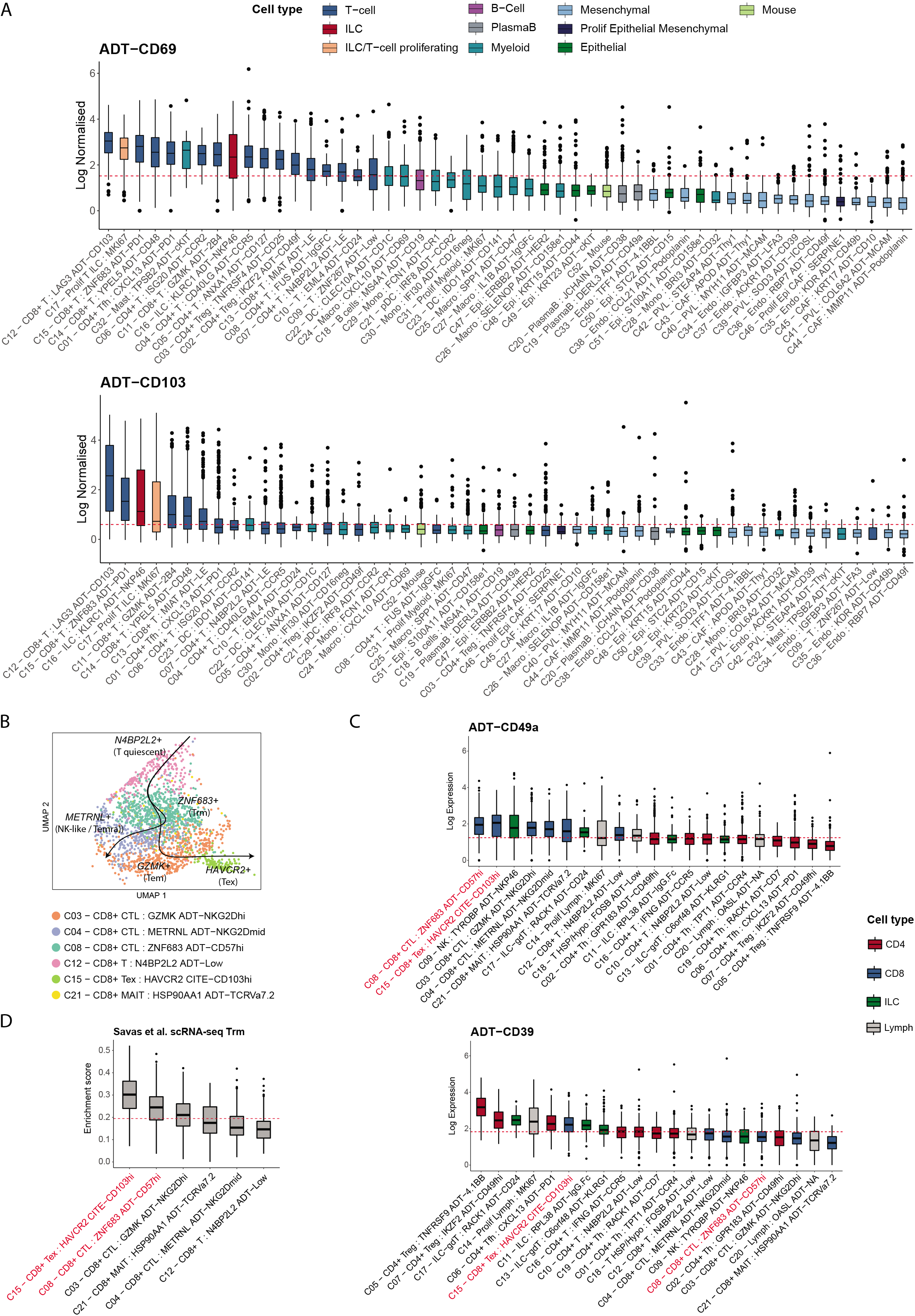
The expression of tissue resident and exhausted T-cell Protein markers and supplementary information for CD8+ T-cell phenotyping. **(A)** Log normalised expression of CD69 and CD103, sorted from high to low (left to right). Bar plots are coloured by broad lineage. Red line marks the median value of each respective feature across all clusters. Boxplots middle line marks the median value, the lower and upper hinges mark the 25% and 75% quantile, the whiskers correspond to the 1.5 times the interquartile range, the black dot’s mark outliers. **(B)** Pseudotemporal analysis of CD8+ T-cells using slingshot. Analysis was performed on RNA values only. **(C)** Log normalised expression of ADT markers CD49a and CD39 across T-cell and ILC clusters, sorted from high to low (left to right). Box plots are coloured by lymphocyte lineage. Red line indicates the median expression for each marker across clusters. Boxplots middle line marks the median value, the lower and upper hinges mark the 25% and 75% quantile, the whiskers correspond to the 1.5 times the interquartile range, the black dot’s mark outliers. **(D)** AUCell enrichment score of signatures derived from Savas et al. (Peter Savas et al., 2018) human breast cancer CD8+ Tissue resident memory T cells (Trm). Red line indicates the median signature score value across all clusters. Boxplots are sorted from high to low (left to right) for each cluster. Boxplots middle line marks the median value, the lower and upper hinges mark the 25% and 75% quantile, the whiskers correspond to the 1.5 times the interquartile range, the black dot’s mark outliers.

**Supplementary Figure 4:**
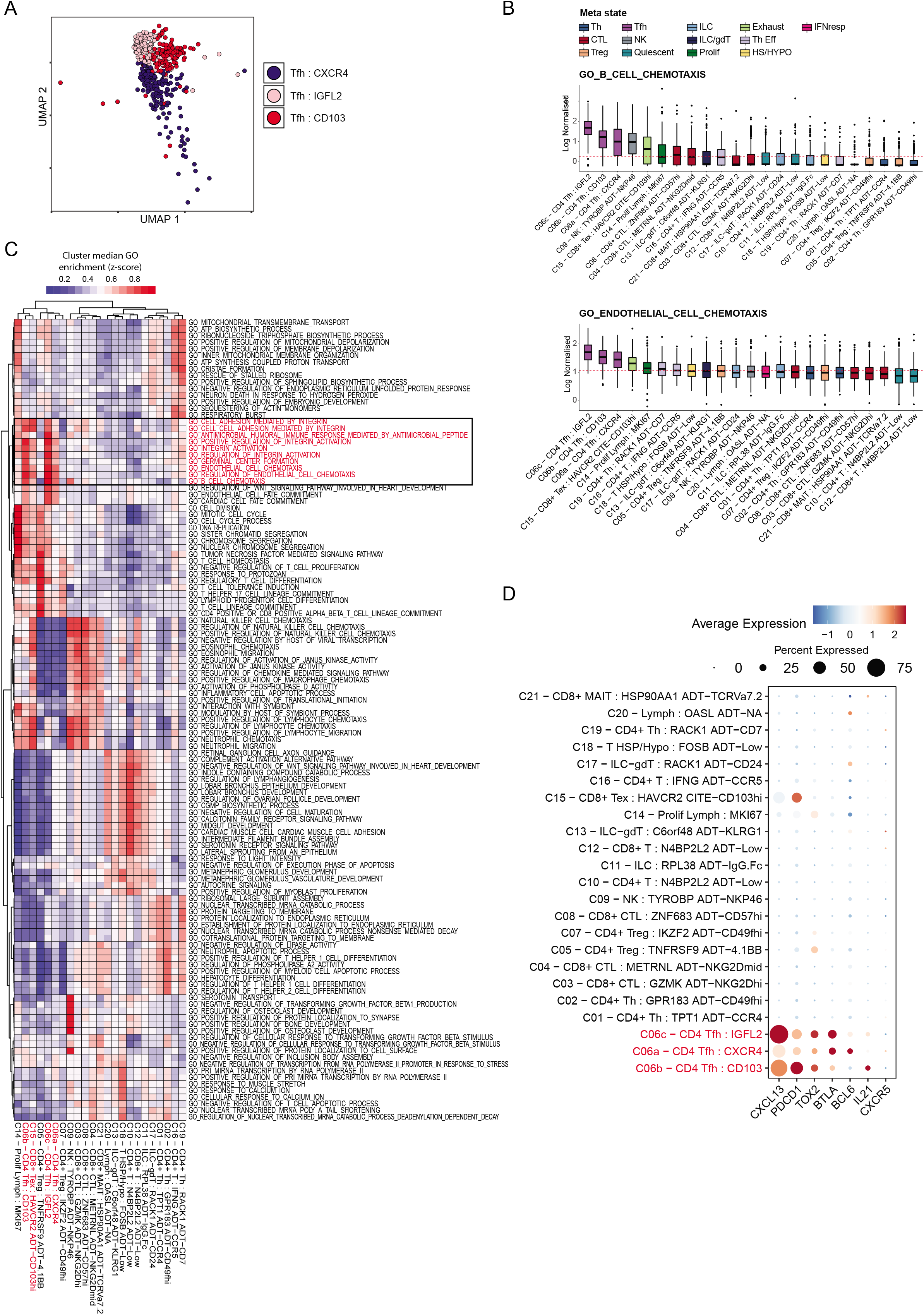
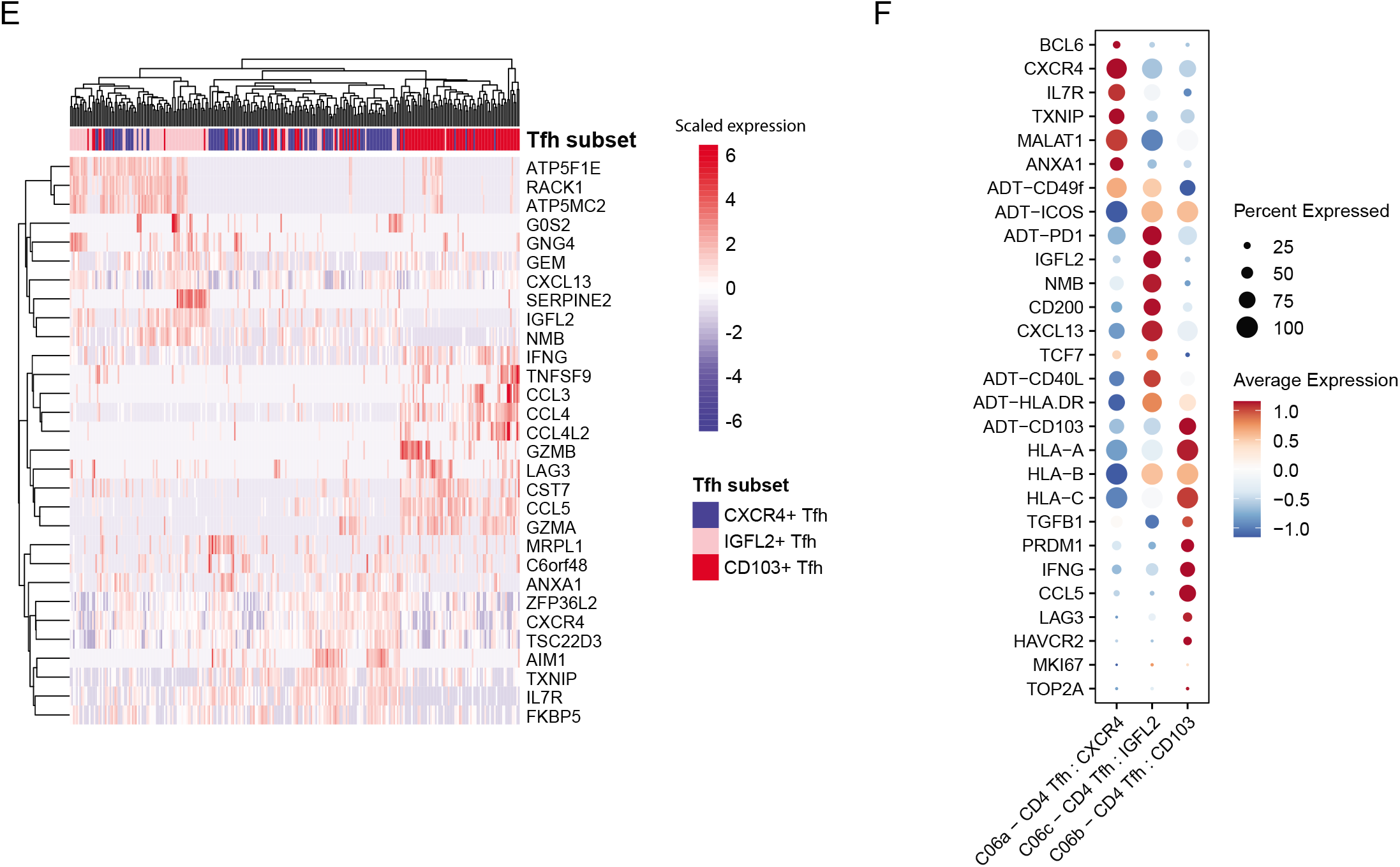
Tfh cells cluster analysis and supplementary phenotyping information of Tfh subsets. **(A)** UMAP of Tfh subsets derived from WNN of ADT and RNA data **(B)** Boxplots of enrichment of GO biological process pathways “GO_GERMINAL_CENTRE_FORMATION”and “GO_B_CELL_CHEMOTAXIS” across T cells & ILCs, sorted from high to low (left to right). Red line indicates the median expression across clusters. Box plots are coloured by lymphocyte lineage. Red line indicates the median expression for each marker across clusters. Boxplots middle line marks the median value, the lower and upper hinges mark the 25% and 75% quantile, the whiskers correspond to the 1.5 times the interquartile range, the black dot’s mark outliers. **(C)** Heatmap of top 5 differentially enriched GO biological process pathways for each cluster based on transcriptome. **(D)** Expression of Tfh cell markers (RNA) across all T-cell and ILC clusters. **(E)** Hierarchical clustering of differentially expressed genes between 3 identified Tfh states. **(F)** Dotplots visualising the expression of z-scaled RNA and ADT markers of interest on a across Tfh subsets.

**Supplementary Figure 5:**
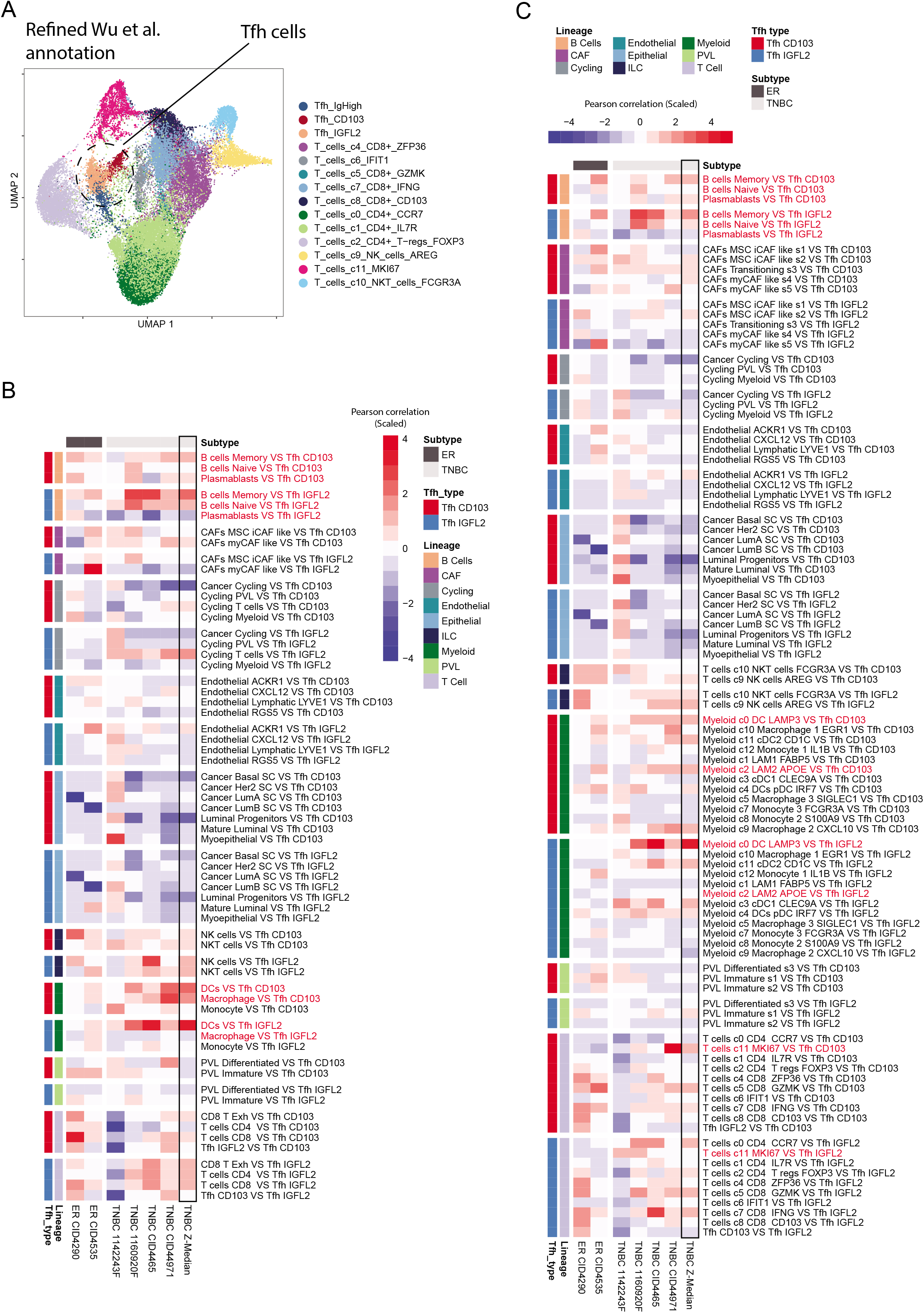
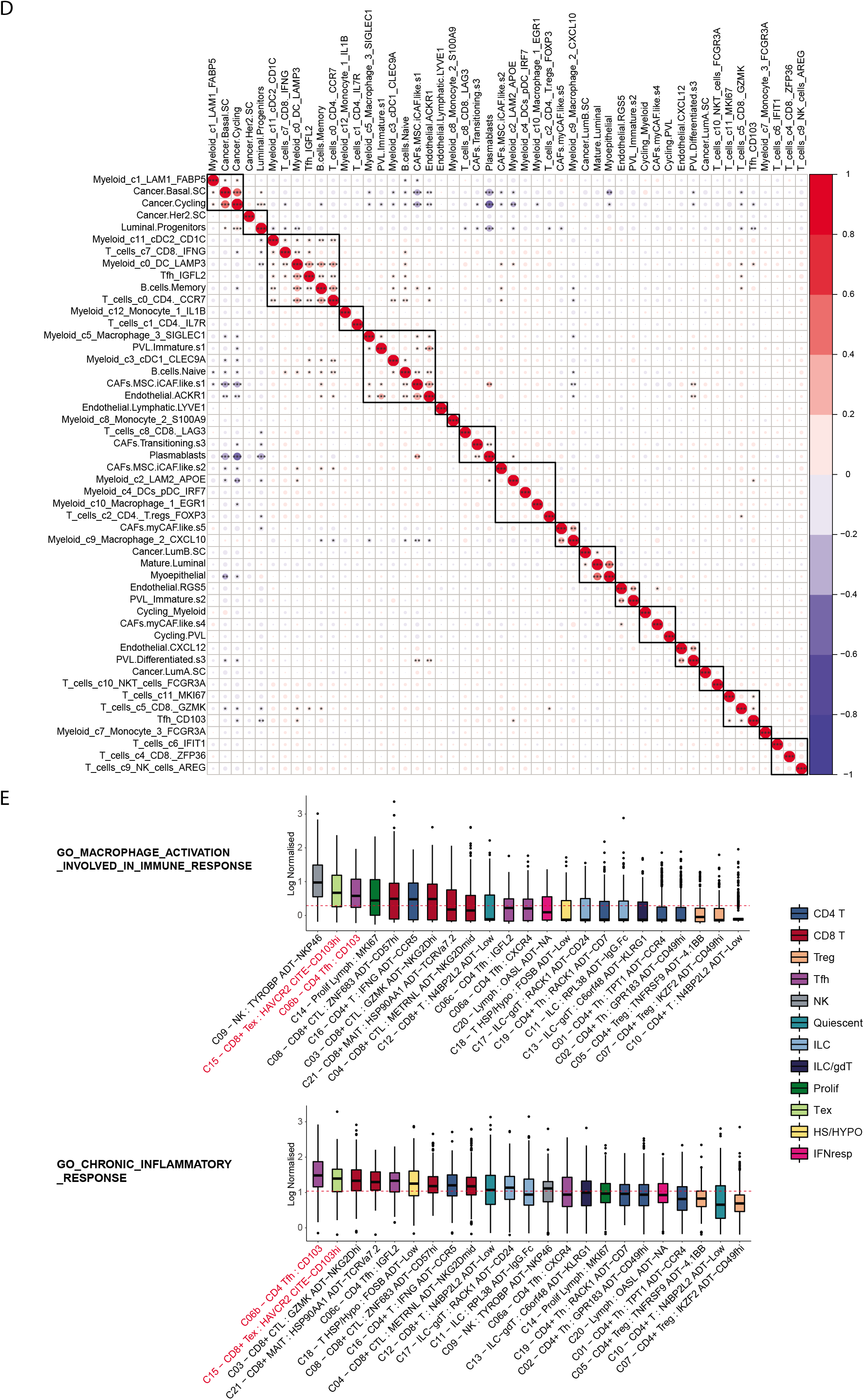
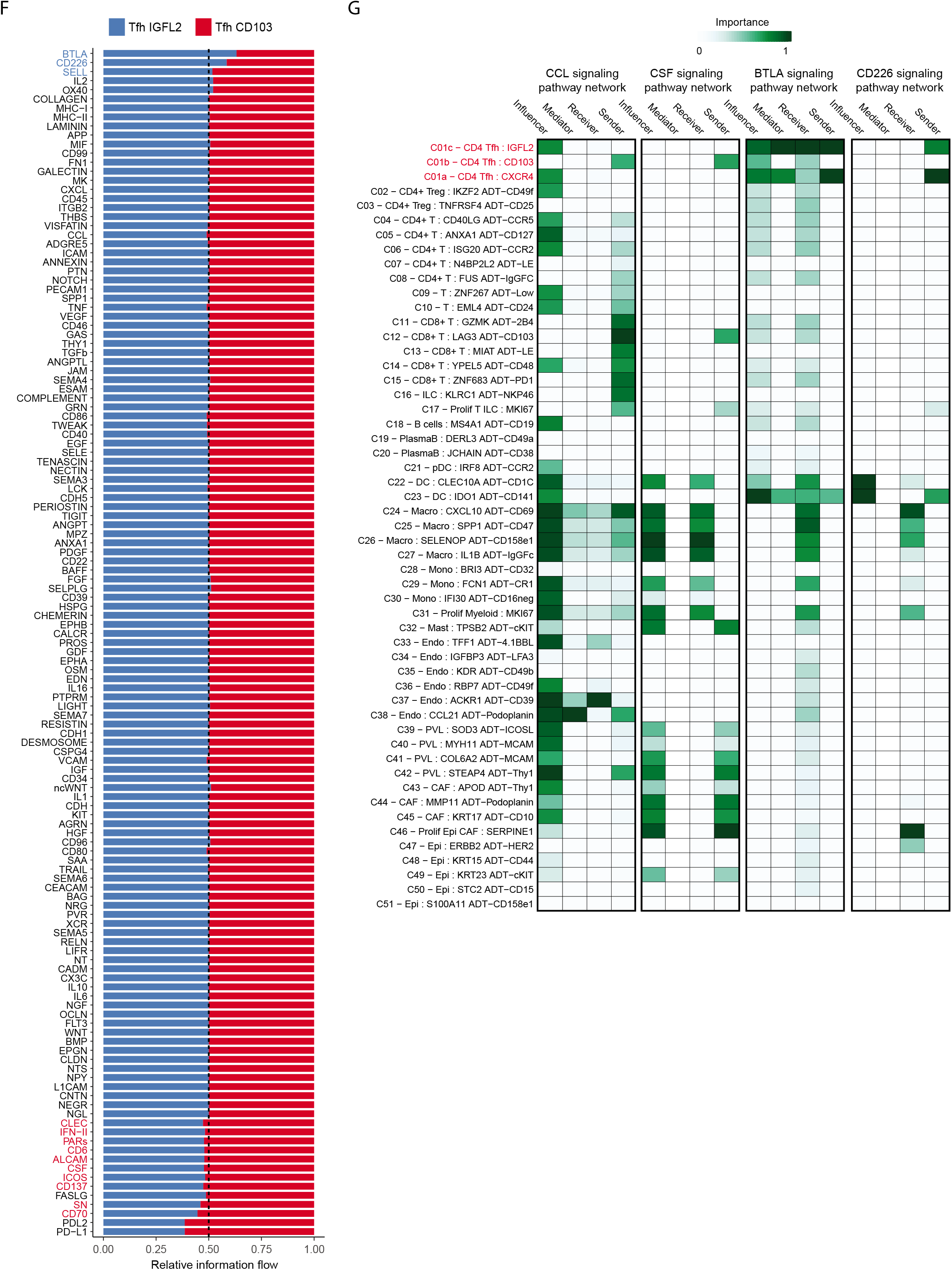
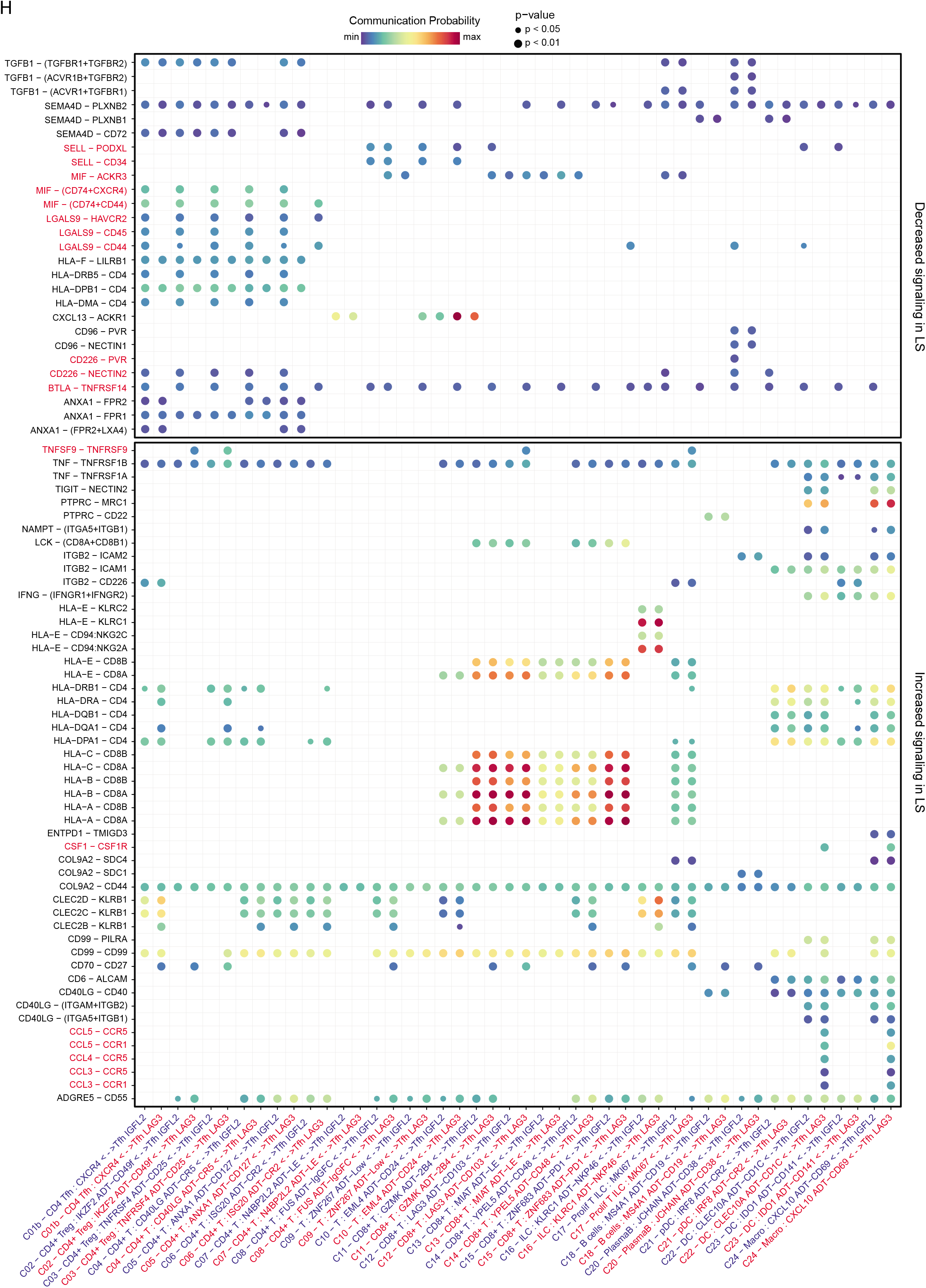
Supplementary data for analysis of Tfh cell subsets spatial co-localisation and cell-cell signalling. **(A)** UMAP of T cells and ILCs from Wu et al. (Wu et al.) with refined annotation of CD4 Tfh cells. **(B)** Pearson correlation heatmap of spatially deconvoluted cell pairs co-localised with CD4 Tfh subsets using Wu et al. data with refined annotations. **(C)** Pearson correlation heatmap of spatially deconvoluted cell pairs co-localised with CD4 Tfh subsets at a more granular resolution. **(D)** Dotplot of Pearson correlation values for spatially deconvoluted cells from (C). **(E)** Gene set enrichment analysis boxplots of GO biological process pathways “GO_B_CELL_CHEMOTAXIS”and “GO_ENDOTHELIAL_CELL_CHEMOTAXIS” across T cells and ILCs, sorted from high to low (left to right). Red line indicates the median expression. Box plots are coloured by lymphocyte lineage. Red line indicates the median expression for each marker across clusters. Boxplots middle line marks the median value, the lower and upper hinges mark the 25% and 75% quantile, the whiskers correspond to the 1.5 times the interquartile range, the black dot’s mark outliers. **(F)** Relative information flow contribution of signalling pathways (the sum of total R-L communication probability found with each signalling pathway) to be differentially enriched in CD4 Tfh cell subsets. Bar plots visualise the weighted variance in contribution when comparing CD103+ Tfh to IGFL2+ Tfh when scaled 0-1. Pathways found to be significant (p < 0.05) are coloured either blue if enriched in IGFL2 Tfh cells, or red if enriched in CD103 Tfh. (G) CCL, CSF and BTLA signalling pathway network information flow across all clusters. ‘Sender’ and ‘Receiver’ reflects direct expression of ligands and receptors (agonistic or antagonistic), ‘Mediator’ and ‘Influencer’ quantifies clusters’ potential role in controlling receptor-ligand expression flow of the pathway within the system (TME). (H) Dotplot visualising receptor-ligand signalling information differentially increased or decreased in expression by CD103 Tfh and IGFL2 Tfh clusters. Red text highlights signalling pathways of interest.

**Supplementary Figure 6:**
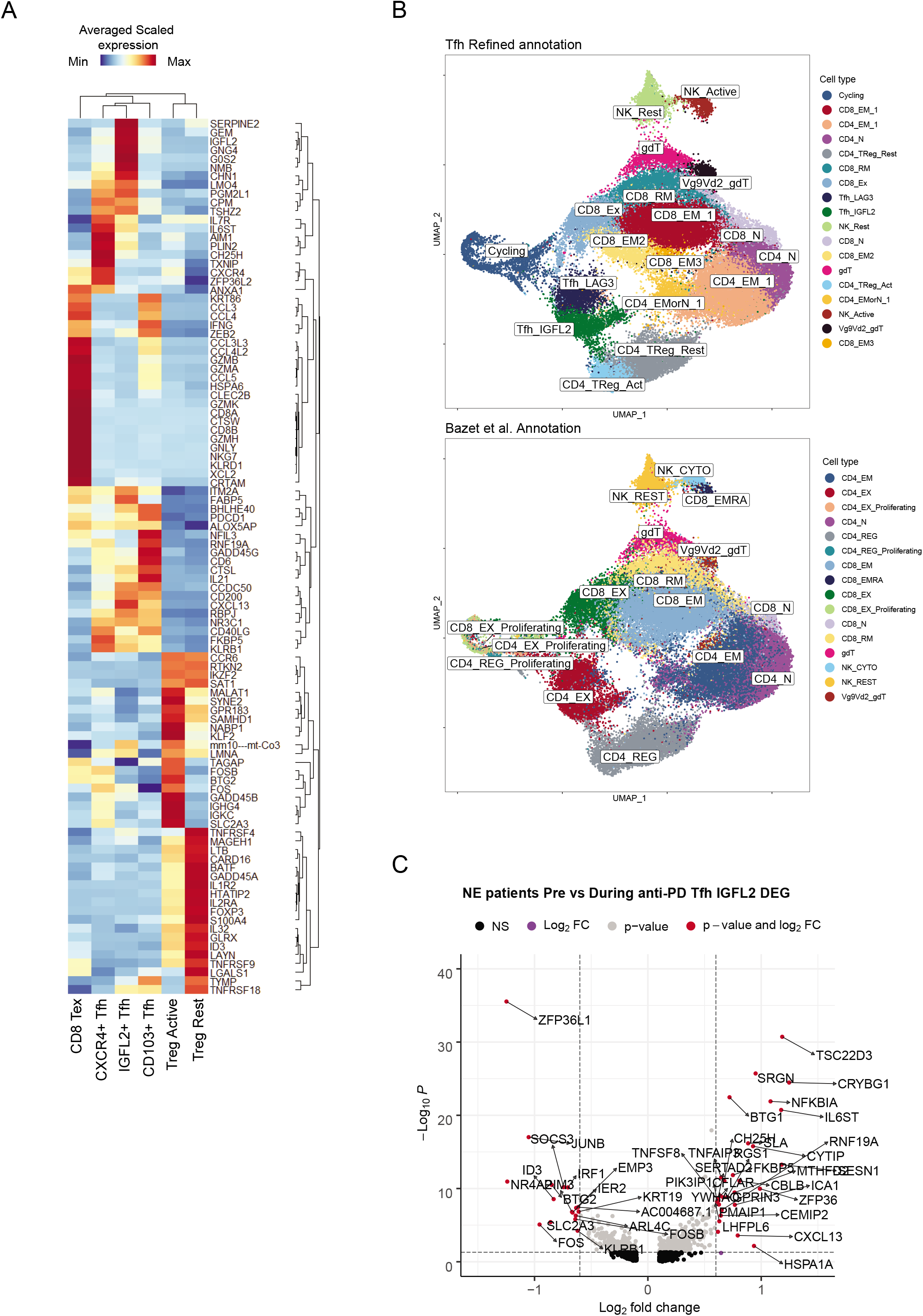
Supplementary data for analysis of Tfh cell subsets clinical relevance. **(A)** Heatmap of top differentiating RNA features from each identified Tfh state in our dataset along with CD8+ Exhausted T cells and Tregs (MAST test; p_adj < 0.05). **(B)** UMAP of Bassez et al. (Ayse Bassez et al., 2021) breast cancer anti-PD-1 cohort T cell dataset. The top plot shows the Tfh redefined annotation used for all downstream analysis in our study. The bottom plot is a UMAP using the Bassez et al. original annotations (Ayse Bassez et al.) **(C)** Volcano plot marking top differentially expressed genes, comparing IGFL2+ Tfh cells of non-expander patients when sampled at baseline vs during treatment.

**Supplementary Figure 7:**
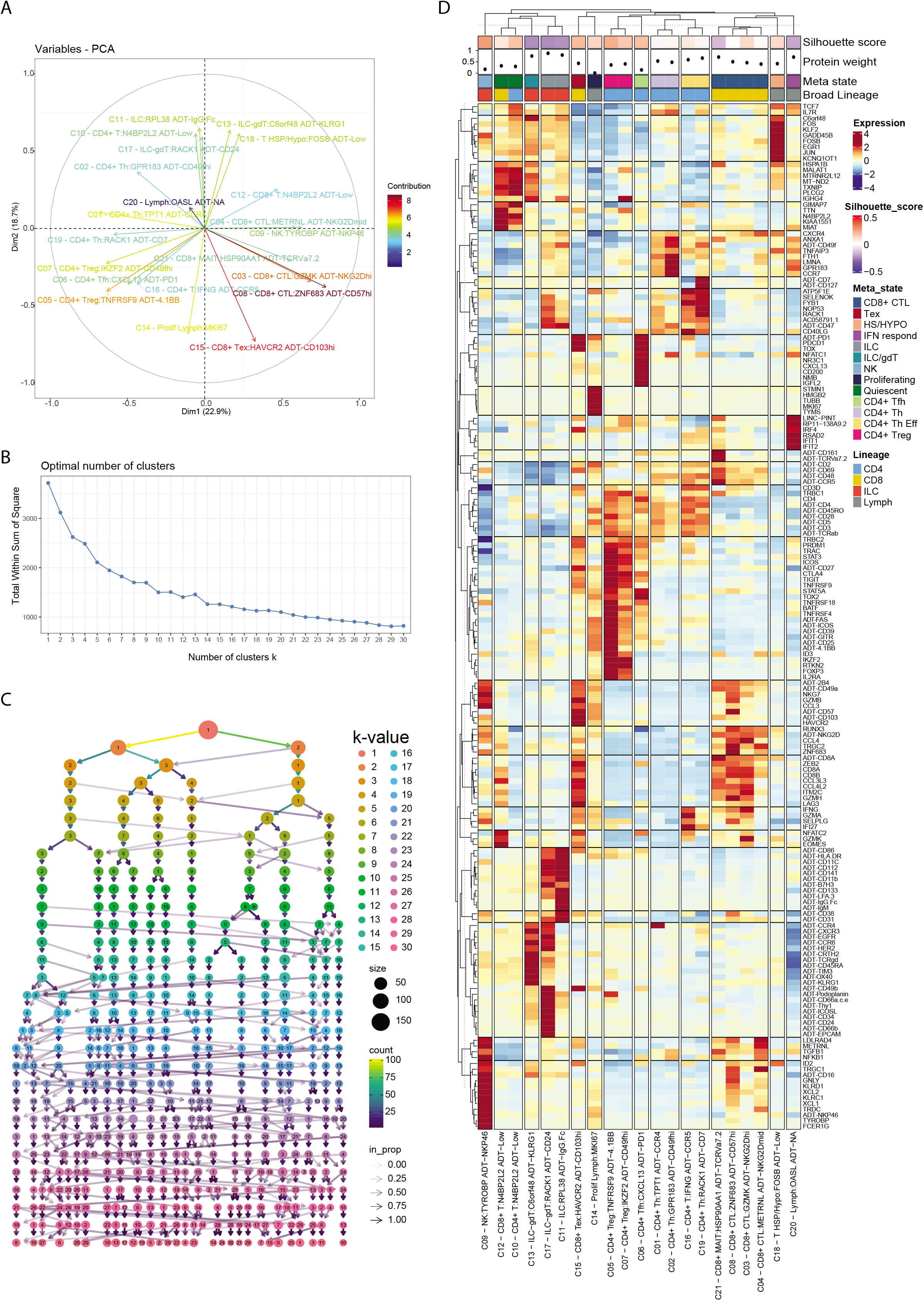
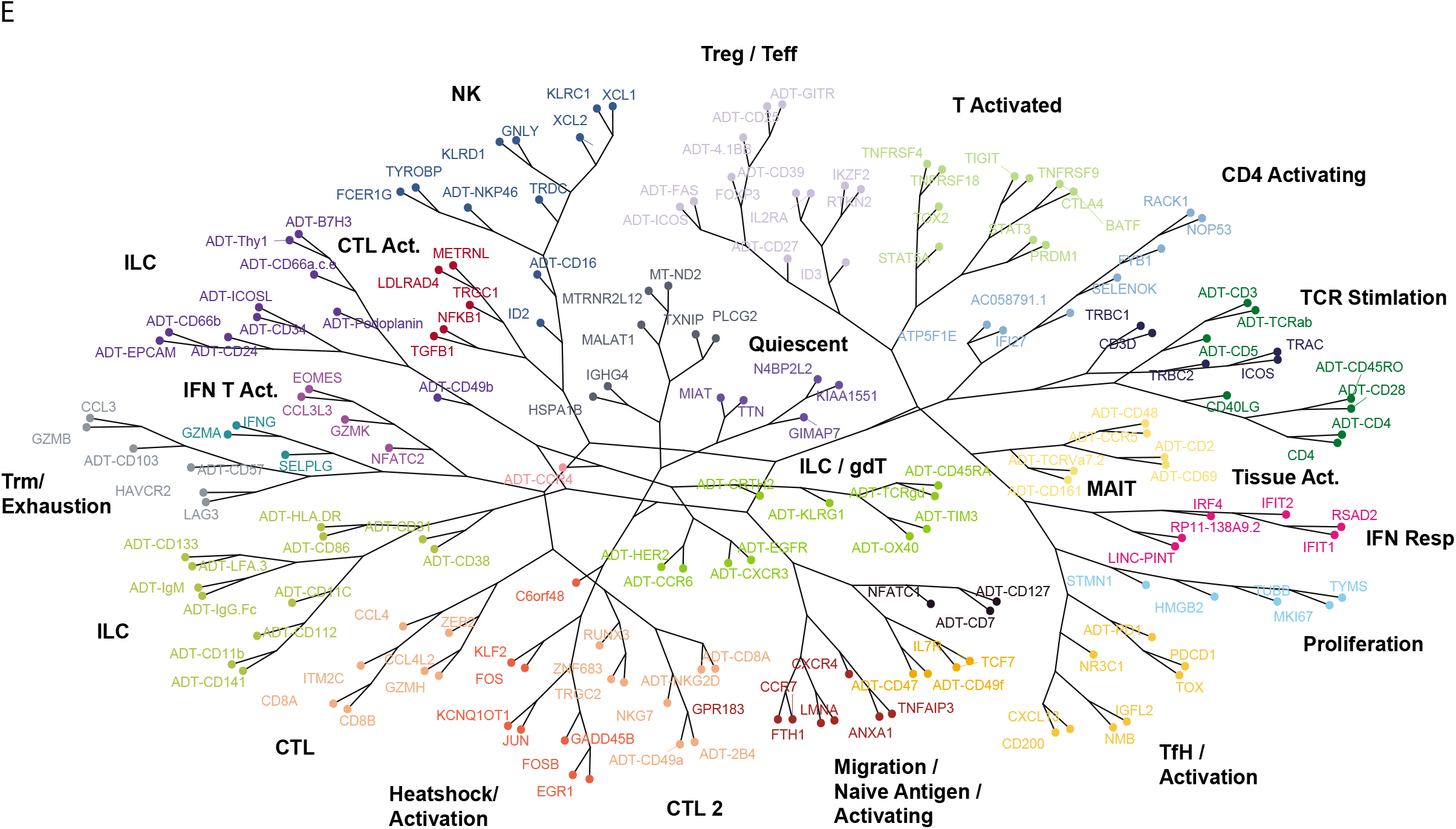
Supplementary data on the co-expression of RNA and ADT features found to drive TIL phenotypes. **(A)** Top PCA of differentially expressed RNA and ADT features, and their contribution to each WNN cluster. **(B)** Scree plot of optimal number of *k* clusters. **(C)** Cluster tree visualising the k-means partitions formed, and their relationship, for every increase in *k* clusters **(D)** Heatmap of top differentiating RNA and ADT features, z-normalised to −4 to +4. Top annotation bar corresponds with, from top to bottom: Median silhouette score of cluster, median protein weighting of cluster, k-means assigned meta state and broad lineage. **(E)** Phylogenetic tree projecting the correlation/co-expression distance of top differentiating RNA and ADT features, irrespective of cluster association. Branch colouring depicts grouping derived from calculated *k* clusters.

